# Concomitant investigation of crustacean amphipods lipidome and metabolome during molting stage by Zeno SWATH Data-Independent Acquisition coupled with Electron Activated Dissociation and machine learning

**DOI:** 10.1101/2023.12.04.569856

**Authors:** Thomas Alexandre Brunet, Yohann Clément, Valentina Calabrese, Jérôme Lemoine, Olivier Geffard, Arnaud Chaumot, Davide Degli-Esposti, Arnaud Salvador, Sophie Ayciriex

## Abstract

**Background:** DIA (Data-Independent Acquisition) is a powerful technique in Liquid Chromatography coupled with high-resolution tandem Mass Spectrometry (LC-MS/MS) in metabolomics and lipidomics, for comprehensive profiling of molecules in biological samples. It provides unbiased coverage, improved reproducibility, and quantitative accuracy compared to Data-Dependent Acquisition (DDA). Combined with the Zeno trap and Electron-Activated Dissociation (EAD), DIA enhances data quality and structural elucidation compared to conventional fragmentation under CID. These tools were applied to study the lipidome and metabolome of the freshwater amphipod *Gammarus fossarum*, successfully discriminating stages and highlighting significant biological features. Despite being underused, DIA, along with the Zeno trap and EAD, holds great potential for advancing research in the omics field.

**Results:** DIA combined with the Zeno trap enhances detection reproducibility compared to conventional DDA, improving fragmentation spectra quality and putative identifications. LC coupled with Zeno-SWATH-DIA methods were used to study molecular changes in reproductive cycle of female gammarids. Multivariate data analysis including Principal Component Analysis and Partial Least Square Discriminant Analysis successfully identified statistically significant features responsible for group segregation. EAD fragmentation helped to identify unknown features and to confirm their molecular structure using fragmentation spectra DB annotation or machine learning. EAD database matching accurately annotated five glycerophospholipids, including the position of double bonds on fatty acid chain moieties. SIRIUS database predicted structures of unknown features based on experimental fragmentation spectra to compensate for database incompleteness.

**Significance:** Reproducible detection of features and confident identification of putative compounds are pivotal stages within analytical pipelines. The DIA approach combined with Zeno pulsing and EAD synergistically enhances detection sensitivity while providing orthogonal fragmentation information, thereby facilitating a comprehensive and insightful exploration of pertinent biological molecules associated with the reproductive cycle of gammarids. The developed methodology holds great promises for identifying biomarkers of significance in the field of environmental assessment.

**Highlights:**

- Advancements in Data-Independent-Acquisition methods for lipidomics and metabolomics.
- Zeno pulsing improves sensitivity in fragment detection and compound identification.
- Comprehensive structure elucidation of lipids and metabolites using EAD fragmentation.

## 1. Introduction

Mass spectrometry (MS)-based omics technologies have revolutionized the ability to measure molecular changes across various levels of an organism’s biological organization, providing a deeper understanding of the impact of stressors on organisms. When coupled with separative techniques such as Liquid Chromatography (LC), the combination enables the separation and detection of numerous molecules from complex biological samples in one run. The comprehensive study of small biomolecules such as lipids and metabolites, known as lipidomics and metabolomics respectively, is needed to unravel the role of these compounds in metabolism and metabolic-related diseases [1,2]. In the last decade, substantial advances have been made in small molecule biomarker discovery, thanks to the ongoing improvement of mass spectrometry (MS) techniques. Advancements were observed in experimental design, data acquisition methods, data preprocessing software, databases, and data processing [3]. Research studies that aim to generate novel data for hypothesis generation of potential biomarkers typically utilize high-resolution mass spectrometry (HRMS) for untargeted data acquisition.

In the untargeted approach, the simultaneous acquisition of high-resolution MS^1^ and the corresponding MS/MS spectra of precursor ions can be achieved using either Data-Dependent Acquisition (DDA) or, conversely, Data-Independent Acquisition (DIA). Indeed, these acquisition modes are nowadays, routinely used to acquire high-resolution fragmentation spectra for thousands of compounds in one single analysis [4]. In DDA, a limited number of precursor ions is selected for fragmentation based on their intensity exceeding a predefined threshold. On the opposite, in DIA, indiscriminate fragmentation of all precursor ions occurs, generating fragment spectra for each individual precursor. While DDA ensures selectivity, it can result in a substantial number of missing values, primarily due to semi-stochastic events in precursor selection for fragmentation. This limitation hampers high-throughput studies. Furthermore, due to the fragmented nature of the acquisition spectra, quantification based on a chromatogram can only rely on MS^1^ data. To address the analytical challenges generated by DDA, the DIA mode exemplified by SWATH (Sequential Window Acquisition of all Theoretical mass spectra) initially emerged as a promising approach in proteomics studies and subsequently expanded its application to various omics fields, especially in the analysis of small molecules [5,6]. In SWATH, the mass range of interest is split into a series of overlapping windows typically ranging from 1 to 25 Da wide. SWATH guarantees comprehensive acquisition of a wide range of compounds in complex biological samples, as each precursor is assigned to a reconstructed fragmentation spectrum. While SWATH has demonstrated its efficacy in proteomics [7–9], it has also been recently explored in the field of metabolomics [10–12] and applied to lipidomics [13,14].

For compound identification, high-resolution tandem mass spectra (MS^2^) with open-access spectral library and related computational tools are needed such as Mass bank of North America (MONA), METLIN, mzCloud, GNPS, SIRIUS and CSI:FingerID, MS-FINDER integrated in MS-DIAL, FBMN, MetGEM [15–27]. All of these tools are primarily developed based on spectra recorded through collision-induced dissociation (CID) fragmentation from HRMS instruments. Recently, a new MS/MS fragmentation mode, Electron Activated Dissociation (EAD) formerly known as Electron-Impact Excitation of Ions from Organics (EIEIO), has been implemented on the latest generation of Q-ToF instrument (ZenoTOF 7600 system from Sciex) [20]. Low-kinetic electron beam irradiation of charged ions generates diagnostic product ions, providing additional structural information not observed in CID spectra. The EAD dissociation mechanisms include electron detachment, internal energy redistribution, and bond cleavage. EAD offers in-depth structural information into the molecular structure, functional groups, double bond positions and stereochemistry [28]. The benefits of EAD have been demonstrated in the identification of lipid species showing its ability to fully characterize a species from a single fragmentation spectrum [29–31]and in metabolomics through the elucidation of cathinone positional isomers [32–34].

On the same instrument, a short linear ion trap located at the end of the collision cell, named the Zeno trap, has been implemented, improving significantly the detection of fragments [35,36]. In the Zeno trap, ions are trapped in an axial pseudopotential well created by applying additional radiofrequency (AC) voltage to all four rods of the trap with the same amplitude and phase [35,36]. The AC field creates a mass-dependent pseudopotential, and by gradually reducing the AC voltage, ions can be sequentially released from the trap, starting from high *m/z* to low, while all ions gain the same kinetic energy. As the ions enter the ToF accelerator, ions with lower *m/z* values catch up with heavier ions. With the trapping and release mechanism of the Zeno trap, a duty cycle greater than 95% can be achieved over a mass range of *m/z* 120 to 2000, resulting in a 5-to-20-fold increase in sensitivity. This capability is particularly valuable for the sensitive and comprehensive acquisition of fragment spectra, aiming to obtain the most accurate and reliable molecular fingerprint of unknown compounds.

Overall, the challenge in global approach applied to lipidomics or metabolomics or the combination of both omics (multi-omics) is the development of robust analytical methodologies for data acquisition, data processing and annotation in routine studies.

In the present work, our aim was to explore the advantages of using LC-Zeno-SWATH-DIA in comparison with the conventional DDA method to improve metabolites and lipids identification. Indeed, after a biphasic extraction protocol (MTBE/MeOH), we examined the lipidome (RPLC fraction) and metabolome (HILIC) of female *Gammarus fossarum,* a freshwater amphipod at two distinct reproductive stages (C1 and D1). We compared data sets acquired in DDA, SWATH-DIA in terms of reproducibly detected features. Next, we evaluated the benefits of the Zeno trapping on the intensity of the fragments generated in SWATH-DIA as well as their numbers and especially for low abundant compounds. We examined the lipidome by RPLC separation (apolar fraction) and the metabolome by HILIC (polar fraction) of female *Gammarus fossarum*, a freshwater amphipod routinely used in Ecotoxicology [37–39], at two distinct reproductive stages (C1 and D1). Multivariate data analysis encompassing an unsupervised approach (principal component analysis, PCA) and supervised modelling (partial least squares discriminant analysis, PLS-DA) was applied to visualize and explain data variability and to highlight specific molecular features involved in reproductive stage differentiation. EAD was then used on the most significant features in the RPLC and HILIC datasets by a high-resolution targeted approach (ZENO-MRM^HR^) to improve structural elucidation, whether using the EAD MS/MS database or machine learning tools such as CSI:FingerID from SIRIUS.

## 2. Materials and methods

### 2.1. Chemicals and reagents

Water, acetonitrile (ACN), methanol (MeOH) and isopropanol (IPA) solvents were obtained from Fisher Scientific (LC-MS grade, Strasbourg, France). Ammonium formate, dichloromethane (DCM), tert-Butyl methyl ether (MTBE), formic acid and ammonium bicarbonate were obtained from Sigma Aldrich (St Quentin-Fallavier, France). Splash® lipidomix® mass spec standard, 1,2-diundecanoyl-sn-glycero-3-phosphocholine (11:0/11:0 PC), 1-tridecanoyl-sn-glycero-3-phosphocholine (LPC 13:0), 1-dodecanoyl-2-tridecanoyl-sn-glycero-3-phosphoethanolamine (12:0/13:0 PE), 1-dodecanoyl-2-tridecanoyl-sn-glycero-3-phospho-(1′-rac-glycerol) (12:0/13:0 PG), 1-dodecanoyl-2-tridecanoyl-sn-glycero-3-phospho-l-serine (12:0/13:0 PS), 1-heptadecanoyl-2-(9Z-tetradecenoyl)-sn-glycero-3-phospho-(1′-myo-inositol) (17:0/14:1 PI), 1,3-diheptadecanoyl-glycerol (d5) (17:0/17:0-d5 DG), N-lauroyl-D-erythro-sphingosine (Cer d18:1/12:0), N-oleoyl(d9)-D-erythro-sphingosylphosphorylcholine (SM d18:1/18:1-d9), 1,3(d5)-diheptadecanoyl-2-(10Z-heptadecenoyl)-glycerol (17:0/17:1/17:0-d5 TG), 1,3(d5)-dipentadecanoyl-2-(9Z-octadecenoyl)-glycerol (15:0/18:1/15:0-d5 TG) were obtained from Sigma Aldrich (St Quentin-Fallavier, France). QReSS kit containing labelled metabolites (stable isotope-labelled molecules was obtained from Cambridge Isotope Laboratories Inc. (Tewksbury, Massachusetts, USA).

### 2.2. Biological sample collection

The collection and maintenance of *G. fossarum* organisms for this study were conducted following previously published protocols [39–42]. Organisms were collected through kick sampling using a net and were selected based on their size, using a series of sieves with a length tolerance of approximately ±1 cm. Upon collection, the organisms were promptly transported to the laboratory and placed in tanks supplied with continuously drilled groundwater under constant aeration. No additional food supply was provided during the study. The temperature of the tanks was maintained at 16±1°C. Female gammarids at specific reproductive stages were identified and sorted (n=12 in C1 stage and n=12 in D1 stage). All collected gammarids were thoroughly washed with deionized water, weighed, and immediately snap-frozen in liquid nitrogen. The frozen samples were then stored at −80°C until further use.

### 2.3. Biphasic extraction procedure for lipids and polar metabolites

The gammarids were individually transferred into homogenization tubes (Precellys® lysing kit CK28R, Bertin Technologies) and homogenized using a Precellys® bead-beating homogenizer (Bertin Technologies) in 300 µL of 150 mM ammonium bicarbonate solution. The homogenization process consisted of two cycles, each lasting 25 seconds at a speed of 500 rpm, with a 10-second pause between cycles. The homogenate was subjected to centrifugation at 13,000 g for 15 minutes, and 250 µL of the resulting supernatant was sampled. Prior to extraction, protein determination assay was performed on the homogenates using the bicinchoninic acid assay. To 200 µL of the homogenate (concentration around 9.4 µg/µL), 226 µL of cold MeOH, 4 µL of the QReSS kit, and 10 µL of Splash® lipidomix® were added. The mixture was then incubated at 4°C on a Thermomixer (Eppendorf AG, Hamburg, Germany) for 1 hour. Following this, 800 µL of cold MTBE was added to the mixture, and samples were mixed at 4°C for 10 minutes before centrifugation at 10,000 g for 5 minutes at 4°C [43]. 880 µL of the upper organic phase, was collected and subsequently evaporated under a nitrogen flow. The resulting dry lipid extracts were resuspended in 15 µL of DCM/MeOH (1:1, *v/v*) and then with 120 µL of IPA/ACN/H_2_O (2:1:1, *v/v/v*). For metabolite analysis, 200 µL of the aqueous lower phase was retrieved, dried under a nitrogen flow, and resuspended in 20 µL of H_2_O/ACN (10:90, *v/v*) to achieve a 10-fold concentration.

### 2.4. QC sample preparation

40 µL of each gammarid homogenate was combined and diluted by a factor of 2 to create a pooled sample that would serve as a quality control (QC) during the LC-MS analysis sequence. From this QC sample, 300 µL was divided into 6 different tubes. To each tube, 315 µL of cold MeOH, 6 µL of the QReSS kit, and 15 µL of Splash® lipidomix® were added. The samples were mixed on an Eppendorf Thermomixer® (Eppendorf AG, Hamburg, Germany) for 1 hour at 4°C. Following this, 1200 µL of cold MTBE was added, and the samples were agitated for 10 minutes at 4°C. After centrifugation at 10,000 g for 5 minutes, 1260 µL of the upper organic phase was collected. The collected phase was then dried under a nitrogen flow, resuspended in 20 µL of DCM:MeOH (1:1, *v/v*), and subsequently with 180 µL of IPA:ACN:H2O (2:1:1, *v/v/v*). For the QC samples of the polar fraction (polar metabolites), the aqueous phases were combined to reach a final volume of 1800 µL. The combined phase was then dried under a nitrogen flow and resuspended in 180 µL.

### 2.5. Untargeted lipidomics and metabolomics by LC-Zeno-SWATH

#### 2.5. 1. Analysis of the organic phase extract

Chromatographic separation of lipid extracts, QCs and blanks were performed on an XSelect CSH C18 column (100 mm x 1 mm, particle size 3.5 µm) heated to 70°C (Waters, Milford, MA, USA) on an Exion LC system. The LC method was adapted from a previously described method [38]. Elution was performed at a flow rate of 200 µL/min with H_2_O:ACN (60:40, *v/v*) containing 10 mM ammonium formate and 0.1% (*v/v*) formic acid for mobile phase A and IPA:ACN:H_2_O (90:9.5:0.5, *v/v/v*) containing 10 mM ammonium formate and 0.1% (*v/v*) formic acid for mobile phase B, employing a linear gradient from 1% B to 100 % B in 13.6 min. The column was washed at 100% B for 2 min before equilibration to 1% B for 2.4 min. The duration of the gradient was 19 min. The autosampler temperature was set at 6°C. The injection volume was 5 µL. MS analyses were performed on a Sciex ZenoTOF 7600 system (Darmstadt, Germany) equipped with an OptiFlow TurboV ion source and operated in positive and negative ion mode in SWATH mode with Zeno trap activated. The following ion source parameters were as follows: a spray voltage of 5.5 kV, a capillary temperature of 300°C, ion source gas 1 and 2 were both set to 70 psi, curtain gas of 30 psi, and CAD gas of 7 psi, the Declustering Potential (DP) is set at 60 V for positive ion mode and −60 V for negative ion mode.

Zeno-SWATH-DIA method using 100 variable-size windows were developed for each chromatography mode in both polarities resulting in four LC-MS/MS methods. SWATH-DIA methods were developed using TOF MS data and SWATH Variable Window Calculator provided by SCIEX. Collision Energy (CE) of each window is calculated using DCE (Dynamic Collision Energy) with the equation for monocharged species (CE = slope*m/z + intercept, slope = 0.05, intercept = 15). The acquisition settings used in this study were as follows: 100 Q1 windows, with an MS^1^ accumulation time of 100 ms. The TOF MS mass range was set from 200 to 1500 Th, while the TOF MS/MS range was set from 100 to 1300 Th. The MS/MS accumulation time was set to 5 ms. External calibrations for both MS and MS/MS were performed before each run using the automated calibration feature. ESI calibration solution SCIEX X500B was used for both positive and negative acquisition modes. During the sequence, samples were injected randomly using each method, QCs and blank samples were injected continuously to ensure the accuracy and reliability of the analysis.

#### 2. 5. 2. Metabolomics analysis

Metabolites were separated on an Acquity Premier BEH amide column (100 mm x 2.1 mm, particle size 1.7 µm) with an injection volume of 5 µL (Waters, Milford, MA, USA). Elution was performed at a flow rate of 200 µL/min with H_2_O containing 10 mM ammonium formate and 0.15 % (*v/v*) formic acid as mobile phase A and ACN/H_2_O (90:10, *v/v*) containing 10 mM ammonium formate and 0.15 % (*v/v*) formic acid as mobile phase B, employing a linear gradient from 0 % A to 56 % A in 18 min. An initial isocratic step was performed during 6 min before the start of the linear gradient. The column was washed at 56 % A for 4 min before returning to 0 % A for re equilibration during 6.8 min. This resulted in a 35 min analysis time. The oven temperature was set at 40 °C and the autosampler temperature was set at 6°C. The following ion source parameters were as follows: a spray voltage of 5.5 kV, a capillary temperature of 500°C, ion source gas 1 and 2 were both set to 70 psi, curtain gas of 50 psi, and CAD gas of 10 psi, the DP is set at 60 V for positive ion mode and −60 V for negative ion mode. CE of each window is calculated using DCE with the equation for monocharged species (CE = slope*m/z + intercept, slope = 0.05, intercept = 15). The acquisition settings were as follows: number of Q1 windows: 100, MS^1^ accumulation time: 100 ms; TOF MS mass *m/z* range: 50-1000 Th, TOF MS/MS *m/z* range: 100-1300 Th, MS/MS accumulation time: 9 msec. External MS and MS/MS calibrations were performed before each run using the automated calibration feature fitted with ESI calibration solution SCIEX X500B for both positive and negative acquisition mode.

### 2.6. Comparison between IDA, SWATH and Zeno-SWATH acquisition mode

QC samples were used for the comparison of SWATH-DIA acquired data towards information-dependent acquisition/ data-dependent acquisition mode (IDA/DDA) and Zeno-SWATH-DIA based on feature detection and annotation respectively. SWATH-DIA acquisition method was developed using the same parameters as Zeno-SWATH-DIA method with the deactivated Zeno trap. IDA/DDA method was developed based on the same source parameters of SWATH-DIA method selecting the 10 most abundant precursor ions for fragmentation (top-10 data dependent MS/MS) with an intensity threshold of 100 cps and dynamic background subtraction. For lipidomics analysis, the TOF MS/MS accumulation time was set at 80 ms, CE was set at 45V with a Collision Energy Spread (CES) of 15V. For metabolomics analysis, the TOF MS/MS accumulation time was set at 140 ms to reach the desired cycle time, CE was set at 30V and CES at 10V.

### 2.7. Zeno-MRM^HR^-EAD

For the structural elucidation of targeted lipids or metabolites of interest, Zeno-MRM^HR^ methods were developed in positive ion mode using the activated Zeno trap and were applied on QCs. Zeno-CID-MRM^HR^ and Zeno-EAD-MRM^HR^ methods were specifically applied to features that exhibited a Variable Importance in Projection (VIP) score above 1.5. The purpose was to enhance the identification or confirmation of the unknown relevant features. Detailed information regarding the method parameters can be found in the supporting information, specifically in Table 1A and 1B.

**Table 1.** Discriminant features in HILIC (A) and RPLC (B). A MS-DIAL identity number is given for each feature. RT as well as the precursor mass-to-charge ratio are given and form the unique serial number RT_*m/z*. The p-value related to the Wilcoxon Mann Whitney non-parametric test is displayed.

| Dataset | MS-DIAL ID | VIP | RT (min) | Precursor ion *m/z* | Molecular formula | Adduct | ppm difference | Identification if available | p-value |
| --- | --- | --- | --- | --- | --- | --- | --- | --- | --- |
| HILIC(+) | 1126 | 2.17 | 14.84 | 434.1649 | C<sub>14</sub>H<sub>23</sub>N<sub>7</sub>O<sub>9</sub> | [M+H]<sup>+</sup> | 3.1 | 14.8_434.16 | 7.40E-07 |
|  | 330 | 1.86 | 10.29 | 244.0793 | C<sub>8</sub>H<sub>15</sub>NO<sub>6</sub> | [M+Na]<sup>+</sup> | 1.9 | N-acetylglucosamine | 6.81E-03 |
|  | 1217 | 1.83 | 6.45 | 461.2764 | C<sub>24</sub>H<sub>36</sub>N<sub>4</sub>O<sub>5</sub> | [M+H]<sup>+</sup> | 0.06 | 6.4_461.28 | 1.00E-02 |
|  | 230 | 1.81 | 10.28 | 204.0875 | C<sub>8</sub>H<sub>15</sub>NO<sub>6</sub> | [M-H<sub>2</sub>O+H]<sup>+</sup> | 1.4 | 10.3_204.09 | 1.45E-02 |
|  | 225 | 1.68 | 15.67 | 202.0690 | C<sub>6</sub>H<sub>13</sub>NO<sub>5</sub> | [M+Na]<sup>+</sup> | 1.0 | 15.7_202.07 | 4.51E-03 |
|  | 1545 | 1.67 | 13.93 | 573.3248 | C<sub>25</sub>H<sub>44</sub>N<sub>6</sub>O<sub>9</sub> | [M+H]<sup>+</sup> | 0.05 | 13.9_573.32 | 2.91E-03 |
|  | 263 | 1.64 | 5.32 | 217.0679 | C<sub>7</sub>H<sub>14</sub>O<sub>6</sub> | [M+Na]<sup>+</sup> | 4.1 | 5.3_217.07 | 1.00E-02 |
|  | 1487 | 1.59 | 14.76 | 549.2684 | C<sub>23</sub>H<sub>33</sub>N<sub>5</sub>O<sub>8</sub> | [M+CH<sub>3</sub>CN+H]<sup>+</sup> | 2.0 | 14.8_549.27 | 1.83E-03 |
|  | 1156 | 1.56 | 13.96 | 442.2668 | C<sub>20</sub>H<sub>35</sub>N<sub>5</sub>O<sub>6</sub> | [M+H]<sup>+</sup> | 0.43 | 14_442.27 | 3.87E-02 |
|  | 1768 | 1.54 | 12.42 | 690.3202 | C<sub>19</sub>H<sub>41</sub>N<sub>15</sub>O<sub>12</sub> | [M+H<sub>2</sub>O+H]<sup>+</sup> | 6.0 | 12.4_690.32 | 1.45E-02 |
| HILIC(-) | 176 | 1.52 | 11.85 | 220.0829 | C<sub>7</sub>H<sub>8</sub>N<sub>4</sub>O<sub>2</sub> | [M+ACN-H]<sup>-</sup> | 2.5 | 11.9_220.08 | 6.81E-03 |
|  | 857 | 1.23 | 1.92 | 398.1262 | C<sub>18</sub>H<sub>25</sub>NO<sub>7</sub>S | [M-H]<sup>-</sup> | 2.9 | 1.9_398.13 | 1.44E-04 |

**B.**
| Dataset | MS-DIAL ID | VIP | RT (min) | Precursor ion *m/z* | Molecular formula | Adduct | ppm difference | Identification if available | p-value |
| --- | --- | --- | --- | --- | --- | --- | --- | --- | --- |
| RPLC(+) | 232 | 1.98 | 5.43 | 565.4033 | C<sub>40</sub>H<sub>52</sub>O<sub>2</sub> | [M+H]<sup>+</sup> | 2.3 | 5.4_565.4 | 4.96E-05 |
|  | 38 | 1.97 | 4.98 | 284.2942 | C<sub>18</sub>H<sub>37</sub>NO | [M+H]<sup>+</sup> | 4.0 | 5_284.29 | 1.73E-02 |
|  | 460 | 1.82 | 8.07 | 742.5756 | C<sub>42</sub>H<sub>80</sub>NO<sub>7</sub>P | [M+H]<sup>+</sup> | 0.7 | PC_O-34:3 | 4.96E-04 |
|  | 278 | 1.72 | 5.41 | 605.3956 | C<sub>40</sub>H<sub>52</sub>O<sub>2</sub> | [M+H<sub>2</sub>O+Na]<sup>+</sup> | 2.5 | 5.4_605.4 | 2.22E-05 |
|  | 780 | 1.59 | 10.65 | 880.7608 | C<sub>43</sub>H<sub>95</sub>N<sub>13</sub>O<sub>4</sub> | [M+Na]<sup>+</sup> | 9.1 | 10.6_880.76 | 4.49E-02 |
|  | 219 | 1.59 | 9.65 | 550.5092 | C<sub>32</sub>H<sub>60</sub>N<sub>4</sub>O<sub>2</sub> | [M+NH<sub>4</sub>]<sup>+</sup> | 5.7 | 9.7_550.51 | 4.49E-02 |
|  | 367 | 1.58 | 9.52 | 679.4302 | C<sub>38</sub>H<sub>60</sub>N<sub>2</sub>O<sub>7</sub> | [M+Na]<sup>+</sup> | 0.6 | 9.5_679.43 | 4.49E-02 |
|  | 217 | 1.57 | 9.66 | 548.5032 | C<sub>35</sub>H<sub>62</sub>O<sub>3</sub> | [M+NH<sub>4</sub>]<sup>+</sup> | 2.0 | 9.7_548.5 | 4.49E-02 |
|  | 958 | 1.56 | 9.40 | 969.7362 | C<sub>55</sub>H<sub>100</sub>O<sub>13</sub> | [M+H]<sup>+</sup> | 12.4 | 9.4_969.74 | 1.73E-02 |
|  | 392 | 1.56 | 9.91 | 698.2554 | C<sub>34</sub>H<sub>39</sub>N<sub>3</sub>O<sub>13</sub> | [M+H]<sup>+</sup> | 1.1 | 9.9_698.26 | 3.32E-02 |
|  | 263 | 1.55 | 9.38 | 587.5485 | C<sub>36</sub>H<sub>72</sub>N<sub>2</sub>O<sub>2</sub> | [M+Na]<sup>+</sup> | 1.2 | 9.4_587.55 | 3.87E-02 |
|  | 562 | 1.54 | 8.24 | 784.5848 | C<sub>44</sub>H<sub>82</sub>NO<sub>8</sub>P | [M+H]<sup>+</sup> | 1.1 | PC_36:3 | 4.96E-05 |
|  | 559 | 1.52 | 11.67 | 782.7221 | C<sub>48</sub>H<sub>92</sub>O<sub>6</sub> | [M+NH<sub>4</sub>]<sup>+</sup> | 2.2 | TG_45:0 | 3.87E-02 |
|  | 515 | 1.52 | 8.01 | 766.5737 | C<sub>44</sub>H<sub>80</sub>NO<sub>7</sub>P | [M+H]<sup>+</sup> | 1.8 | PC_O-36:5 | 4.51E-03 |
|  | 535 | 1.51 | 9.45 | 774.6373 | C<sub>44</sub>H<sub>88</sub>NO<sub>7</sub>P | [M+H]<sup>+</sup> | 0.5 | PC_O-36:1 | 3.33E-05 |
|  | 471 | 1.5 | 10.18 | 750.4069 | C<sub>30</sub>H<sub>51</sub>N<sub>15</sub>O<sub>8</sub> | [M+H]<sup>+</sup> | 7.3 | 10.2_750.41 | 4.49E-02 |
| RPLC(-) | 130 | 1.45 | 8.27 | 828.5743 | C<sub>44</sub>H<sub>82</sub>NO<sub>8</sub>P | [M+HCOO]<sup>-</sup> | 1.4 | PC_36:3 | 7.20E-05 |
|  | 38 | 1.34 | 3.66 | 327.2326 | C<sub>22</sub>H<sub>34</sub>O<sub>3</sub> | [M-H<sub>2</sub>O-H]<sup>-</sup> | 0.6 | 3.7_327.23 | 3.60E-03 |
|  | 91 | 1.3 | 8.60 | 698.5120 | C<sub>39</sub>H<sub>74</sub>NO<sub>7</sub>P | [M-H]<sup>-</sup> | 0.7 | 8.6_698.51 | 1.40E-05 |
|  | 25 | 1.12 | 3.08 | 301.2170 | C<sub>20</sub>H<sub>32</sub>O<sub>3</sub> | [M-H<sub>2</sub>O-H]<sup>-</sup> | 0.9 | FA_20:5 | 4.50E-03 |

### 2.8. Data processing and preprocessing

Data acquisition was performed in two batches: one for HILIC and one for RPLC, considering both polarities for each batch. The acquired data was then exported and processed using MS-DIAL for peak detection, adduct attribution, MS/MS spectral deconvolution, identification, and alignment. To correct for signal drift, LOESS (Locally Estimated Scatterplot Smoothing) was employed using QCs [44]. In the case of RPLC, isotopically labelled standards lipids were used for normalization, while no labelled metabolites were used in HILIC due to their low signal intensity and poor repeatability, as indicated by the QC Relative Standard Deviation (RSD).

To reduce the total number of variables and eliminate non-relevant features, a set of filters was applied to the data prior to multivariate statistical analysis. Four filters were applied as follows: (i) Blanks: Features with average areas in both sample groups less than 10 times that of the blanks were excluded. (ii) QC: Features with an RSD greater than 30% in the QCs were removed. (iii) Missing values: Features present in less than four samples (considering C1 and D1) were eliminated. (iv) Dead time: Features with a retention time (RT) too close to the dead time were removed.

### 2.9. Statistical analyses

Statistical analyses were carried out using the R program (v4.2.1) with the ropls and mixOmics packages. Initially, Principal Component Analysis (PCA) was performed on all the different datasets. PCA enabled the assessment of analysis quality by projecting blanks and QC samples. It also facilitated the identification of differences between samples (reproductive stages C1 vs D1) and the detection of potential outliers. Subsequently, various models were developed, specifically Partial Least Squares-Discriminant Analysis (PLS-DA), to identify and select the variables responsible for the observed separations between the two reproductive stages. Variable selection was performed based on VIP scores and S-plots. The statistical significance of the models was assessed using the confusion matrix and validation set permutations. The significance of variables was calculated using the Wilcoxon-Mann-Whitney non-parametric univariate test.

### 2.10. Compounds annotation with MS-DIAL module and SIRIUS

The raw data were acquired using SCIEX OS software (v3.0) and subsequently processed with MS-DIAL software (v4.9). The identification of compounds was performed using RIKEN Institute ESI-MS/MS DB from authentic standards databases, based on CID MS/MS spectra. For comprehensive lipid structural identification utilizing EAD fragmentation, the latest version of MS-DIAL (v5.1) was employed, which includes the MS/MS-EIEIO lipidomics database. To predict the structure of unknown metabolites based on CID and EAD spectra, SIRIUS (v5.6.3) in conjunction with CSI:FingerID was utilized to obtain partial information about the molecular structure [21–27].

### 2.11. Data availability

Data will be made available on request.

## 3. Results and discussion

### 3.1. Development of SWATH-DIA method for multi-omics analysis

DIA analysis involves the collection of MS/MS spectra from wide, pre-defined precursor selection windows, typically exceeding 1 Da. In each cycle, an MS^1^ spectrum is often acquired to help in linking the precursors and fragments. The SWATH method utilizes the distribution of wide MS^1^ windows, also exceeding 1 Da, where all precursor ions are indiscriminately fragmented. This results in the acquisition of one MS/MS spectrum per SWATH window, which requires a deconvolution step to correctly associate fragments within each detected precursor ion. The complexity of deconvolution depends heavily on the number of fragmented precursor ions within the window. Consequently, a wide SWATH window positioned in a densely populated precursor area can complicate the assignment of fragments due to the intricacies of the MS/MS spectrum. To alleviate this, variable-sized windows can be employed, with smaller ones designated for dense precursor regions to facilitate deconvolution. To develop a customized SWATH method, two precursor ion scans were performed: one from a blank sample and another from a QC. The blank chromatogram was subtracted from the QC sample chromatogram to ensure that the window distribution was solely based on the precursor ions originating from the gammarid extract. As a result, four SWATH-DIA methods were developed for the characterization of the gammarid lipidome and metabolome: RPLC(+), RPLC(-), HILIC(+), and HILIC(-). The RPLC methods were specifically designed for lipids and nonpolar compounds, considering the LC method and mass ranges, while HILIC focused more on polar metabolites. Each method utilized 100 windows with a minimum width of 3 Da and a window overlap of 1 Da. For RPLC, the mass range was set to 200-1500 *m/z*, while for HILIC, it was set to 50-1000 *m/z* (Fig. S1). The blue curves represent the normalized density of precursor ions across the *m/z* range, demonstrating the separation, ionization, and detection of diverse biomolecule classes. The profiles differ between methods, reflecting the separation of distinct biomolecule classes. The orange curves represent the distribution and width of the windows, where a lower curve indicates a narrower width. It can be observed that the distribution is concentrated in specific areas for RPLC(+) and HILIC(-). These dense areas primarily correspond to the mass range of glycerophospholipids, sphingolipids, and glycerolipids in RPLC (∼*m/z* 600-900), while for HILIC, they correspond to diverse abundant metabolite classes that are sufficiently ionized in ESI(-), such as small organic acids, nucleotides, bile acids, and sugars. The distribution is broader for ESI(-) RPLC and ESI(+) HILIC methods (Fig S1A-D).

### 3.2. Comparison of the IDA/DDA and SWATH-DIA acquisition modes for multi-omics analysis

We hypothesized that the ZENO trap pulsing combined with the SWATH acquisition mode would improve the detection reproducibility and the duty cycle. QC samples were prepared for lipidomics and metabolomics and were injected in triplicate for each chromatography separation and ionization mode (positive and negative). We compared the data obtained from the IDA/DDA mode in TOP10, SWATH-DIA, and SWATH-DIA with Zeno on. First, we compared the DDA and SWATH acquisition modes, considering only the features that were detected and aligned in all three replicates of each acquisition mode for the comparisons. As shown in Fig. 1, SWATH-DIA exhibited a higher number of reproducible detected features compared to DDA, with a 55% increase for RPLC(+) and 58% increase for HILIC(+), as well as a 22% increase for RPLC(-) and 129% increase for HILIC(-). These results suggest a significant improvement in detection reproducibility for SWATH method. Subsequently, to further elucidate the advantages of Zeno pulsing, a comparison was conducted between Zeno-SWATH-DIA and SWATH-DIA (Zeno trap off), focusing on MS/MS spectra and, consequently, putative identifications.

**Figure 1:**
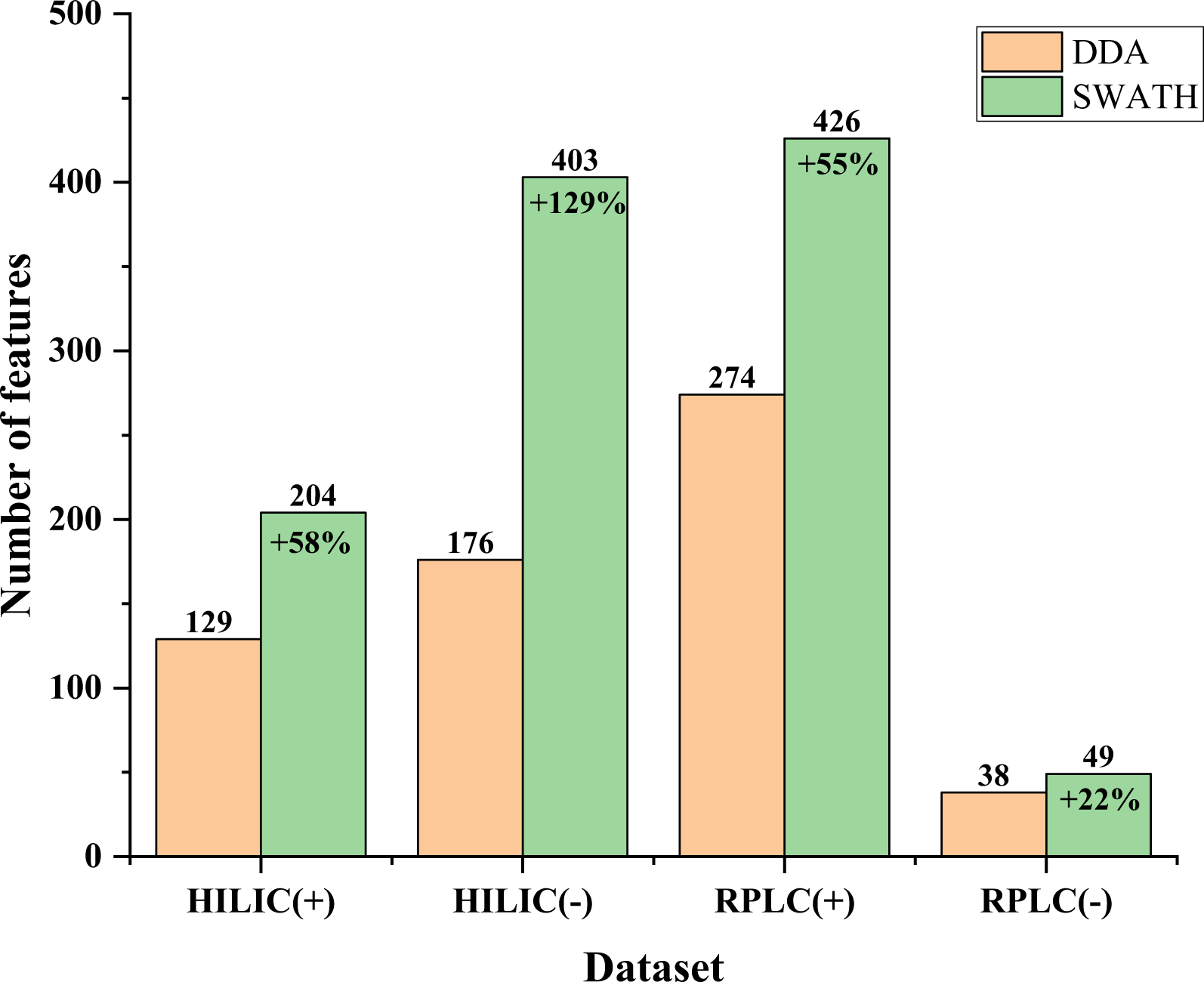
Comparison of SWATH-DIA and DDA (Top 10) for gammarid feature detection in the different LC-MS/MS methods. Features detected in DDA are colored in yellow and features detected in SWATH-DIA are colored in blue. Analyses were performed in triplicate.

### 3.3. Comparison of SWATH-DIA and ZENO-SWATH-DIA

For each dataset, we determined the features shared by SWATH-DIA and Zeno-SWATH-DIA. We then extracted and compared the corresponding MS/MS spectra for each pair of features in terms of fragment intensity for fragments present in both, as well as the total number of fragments for shared features (Fig. 2). In HILIC(+), a total of 343 fragments were used, 523 in HILIC(-), 53 in RPLC(+), and 28 in RPLC(-). To assess the increase in MS/MS sensitivity using the Zeno trap, the comparison was conducted at three n-fold increases: fragments detected with an intensity increase above 5-fold (> 5-fold), above 7-fold (> 7-fold), and more than a 10-fold increase (> 10-fold) (Fig. 2A). Overall, the > 5-fold increase accounted for approximately 50-60% of fragments depending on the dataset, with a combined percentage of 52% (Fig. 2A). The > 7-fold increase covered 40% of the total fragments, while between 19-36% exhibited a > 10-fold increase, resulting in a global percentage of 22%. Notably, higher percentages were observed for RPLC datasets in terms of > 7-fold and > 10-fold increases (Fig. 2A). RPLC emphasizes lipid species with fewer CID fragments compared to metabolites, making the increase more obvious on a smaller number of fragments, thus explaining the higher percentages for RPLC datasets. In conclusion, the Zeno trap significantly enhances MS/MS sensitivity for small biomolecules (lipids and metabolites) in both chromatography modes. This observation is particularly valuable for the molecular characterization of unusual biomolecules. This leads us to question whether the Zeno trap facilitates the detection of initially undetected low-abundance fragments. To demonstrate this, we highlighted the Zeno trap’s impact on the fragment number gain for shared features at three n-fold gains: > 2-fold gain, > 4-fold gain, and > 6-fold gain (Fig. 2B). Globally, out of the 699 total shared features, 78% of them obtained at least twice as many fragments when using the Zeno trap. This trend is consistent across HILIC(-), RPLC(+), and RPLC(-) datasets, with a 32-38% increase for > 4-fold gains and an 11-18% increase for >6-fold gains (Fig. 2B). However, HILIC(+) demonstrated notably high percentages with 67% for > 4-fold gains and 60% for > 6-fold gains (Fig. 2B). This suggests that small positively ionized metabolites may have a significant number of initially undetected fragments due to their low intensity, which the Zeno trap makes visible even in small quantities. This comprehensive fragmentation spectrum is crucial for MS/MS-based identification. To illustrate these observations, we chose four fragmentation spectra from HILIC and two from RPLC to showcase the benefits of the Zeno trap with raw data. These examples align with the aforementioned trends, although in some cases, low-intensity fragments detected without the Zeno trap disappeared when it was enabled (Fig. S2-7). This raises concerns about their reliability and whether they are relevant fragments or artifacts introduced during the deconvolution step. When using Zeno-SWATH, the Zeno trap accumulates fragment ions indiscriminately from a SWATH window before sending them to the detector. Subsequently, algorithmic deconvolution utilizing least square fitting is employed to assign the correct MS/MS chromatogram to precursors based on model peak fitting. The algorithm searches for the fragments most correlated to the precursor by comparing their peak shapes. A strong correlation leads to the fragment being assigned to its corresponding precursor, resulting in a cleaned fragmentation spectrum for each precursor. This deconvolution step is facilitated by the improved fragment intensity, as the algorithm can more reliably fit the peak model when the MS/MS signals are sufficiently intense. This leads to a more consistent combination of fragments and precursors, which may explain both the appearance and disappearance of fragments when using the Zeno trap. Incorrectly assigned fragments are eliminated, while relevant ones that were previously not intense enough are now detected with sufficient intensity for accurate assignment.

**Figure 2:**
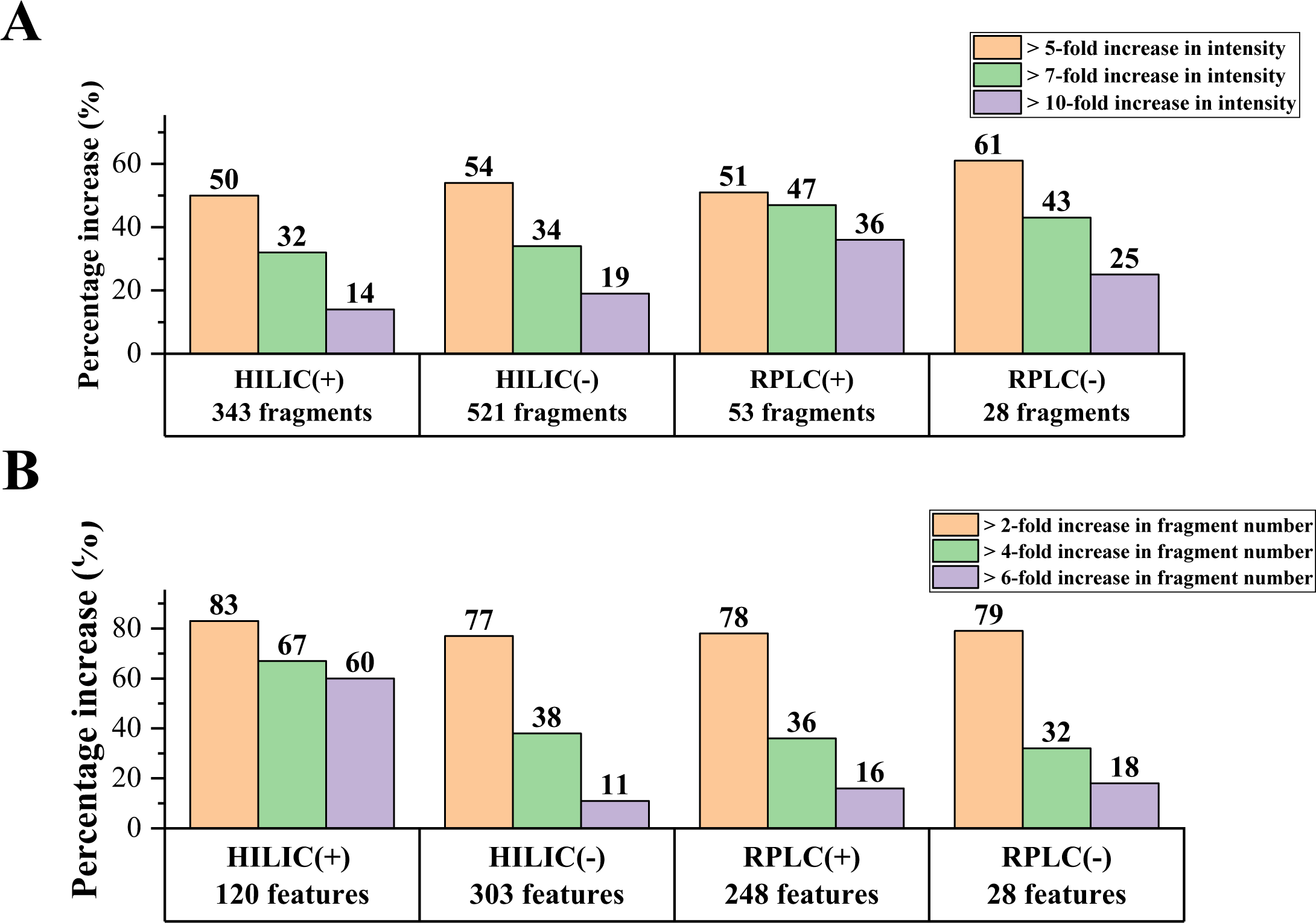
Evaluation of Zeno trap pulsing on fragment intensity (A) and the number of detected fragments (B). The evaluation was performed on the comparison of the intensity of the fragments detected both with Zeno on and Zeno off (A). The evaluation was performed on the comparison of the number of fragments detected both with Zeno on and Zeno off (B).

As mentioned earlier, the Zeno trap’s ability to enhance detection sensitivity and the comprehensiveness of fragmentation spectra represents a significant advantage for MS/MS-based annotation. To illustrate the benefits of the Zeno trap for putative MS/MS identifications, we used a combination of the Reverse Dot Product and the fragment presence ratio of MS-DIAL for comparison. This combination relies solely on the presence of reference fragments in the experimental spectrum to generate an annotation score, without being biased by artifacts. Putative identifications were conducted using the same MS/MS databases and identification score thresholds. The comparison was performed at three progressive identification scores: 50%, 70%, and 80% similarity scores (Fig. 3). In our case study, a score above 50% provides relevant insights into the structure of the unknown feature without necessarily confirming its identification. Scores above 70% indicate highly plausible identity suggestions, while scores above 80% practically confirm the putative identification. In HILIC, an improvement of 41% in identifications with a score above 50% was observed when considering both polarities together (Fig. 3). This improvement increased to 46% for scores above 70% and 43% for scores above 80% (Fig. 3). For RPLC(-), no significant differences were observed due to the limited number of putatively identified features, providing little room for meaningful comparison. However, in RPLC(+), there was a substantial increase of 92% for scores above 70% and 72% for scores above 80% (Fig. 3). Overall, there was a 57% increase in almost certain identifications with a score above 80% when considering both HILIC and RPLC datasets together. Therefore, the Zeno trap demonstrates promising results for the putative identification of unknown small biomolecules in complex biological matrices. It is important to note that MS/MS-based identification heavily relies on the database used meaning that the number of putative identifications is dependent on the exhaustiveness of the database, (the number of spectra recorded), the spectrometer used for the acquisition as well as the experimental conditions such as the acquisition mode.

**Figure 3:**
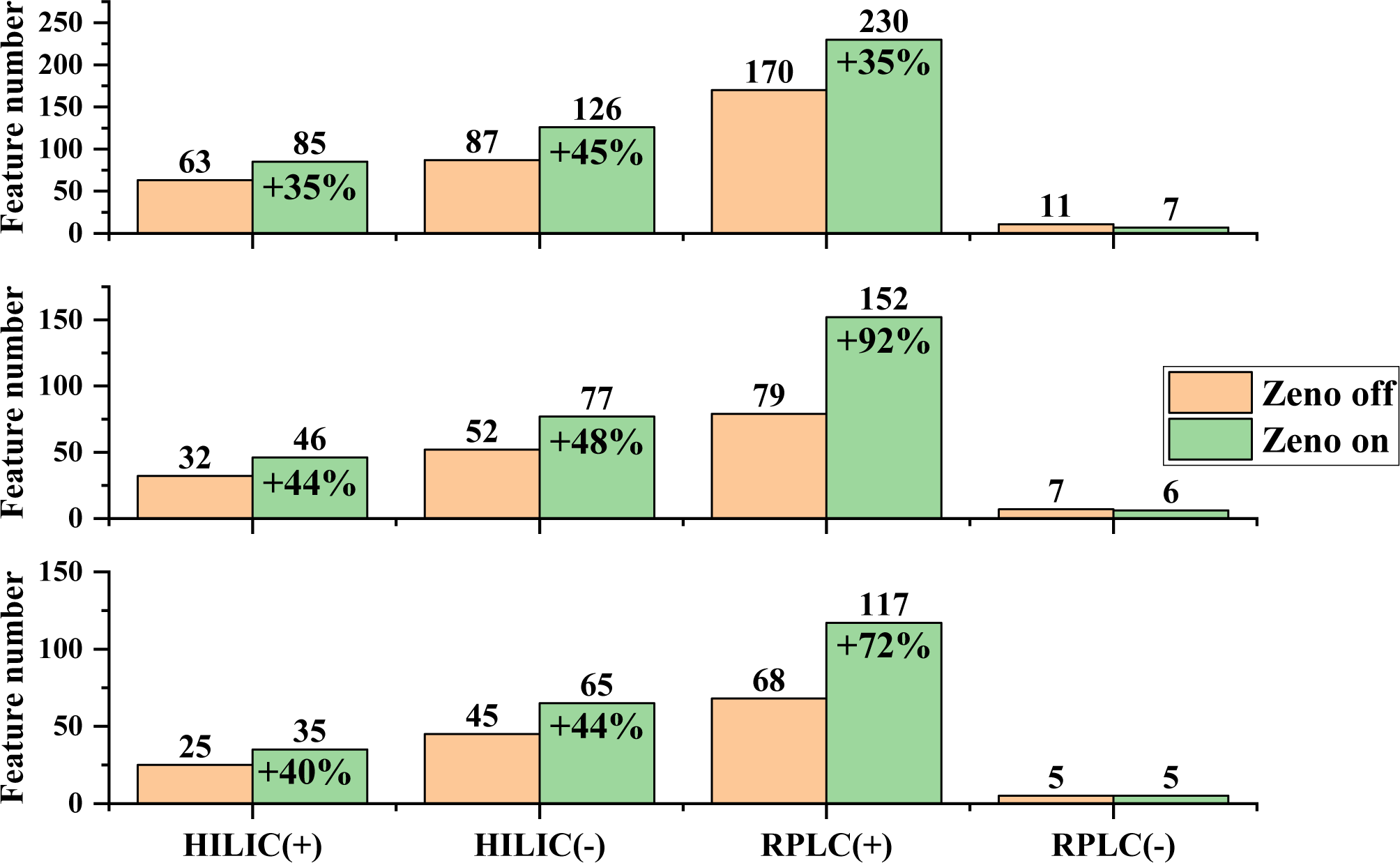
Number of identified features by MS/MS database match with Zeno trap on/off. Putative identification performed by database match at different similarities score: above 50% (A), above 70% (B) and above 80% (C). The yellow bars represent features detected consistently in triplicate without the Zeno trap, while the blue bars represent those detected with the Zeno trap.

### 3.4. Multivariate statistical analysis applied on HILIC and RPLC datasets

To assess the suitability of the Zeno-SWATH-DIA method, we investigated molecular differences of two reproductive stages of female crustacean *Gammarus fossarum*. After a biphasic extraction (MTBE/MeOH), both fraction aqueous (polar metabolites) and organic (nonpolar compounds, mainly lipids) were analysed in HILIC and RPLC respectively. Collected data in both polarities were subjected to unsupervised analysis. PCA analysis of the HILIC(+) dataset exhibited a clear separation between the C1 and D1 groups as well as a well-defined cluster of QCs after correction and a cluster of off-centre blanks (Fig. S8). Similar trends were observed in HILIC(-), RPLC(+), and RPLC(-) data sets (Fig S8). PLS-DA models were built to identify the variables responsible for the separation between C1 and D1 organisms (Fig. 4). The quality of these models was assessed using bootstrap resampling. The R^2^X, R^2^Y, Q^2^Y, and RMSEE values obtained from the PLS-DA models are presented in the supplementary data (Table S4). These values were used to determine the optimal number of principal components (PCs). Evaluation of the model performance using two PCs revealed R^2^Y values ranging from 0.71 to 0.94 and corresponding Q^2^ values ranging from 0.42 to 0.90 (Table S4). Clear separation between the C1 and D1 stages was observed in all four datasets, with the separation being more pronounced and reliable in the HILIC datasets. VIP scores derived from the PLS-DA analysis were used to select the variables responsible for stage separation. VIP scores greater than 1.5 for HILIC(+), 1.2 for HILIC(-), 1.5 for RPLC(+), and 1.1 for RPLC(-) within each dataset were used as selection criteria (listed in Table 1A-B). This criterion resulted in a list of 32 features (each represented by the pair, RT_*m/z*), including 10 features for HILIC(+), 2 for HILIC(-), 16 for RPLC(+), and 4 for RPLC(-). These 32 significant features, along with their MS-DIAL identification number, VIP score, RT, precursor mass, identification (if available), and p-value from the Wilcoxon test, are listed in Table 1A-B.

**Figure 4.**
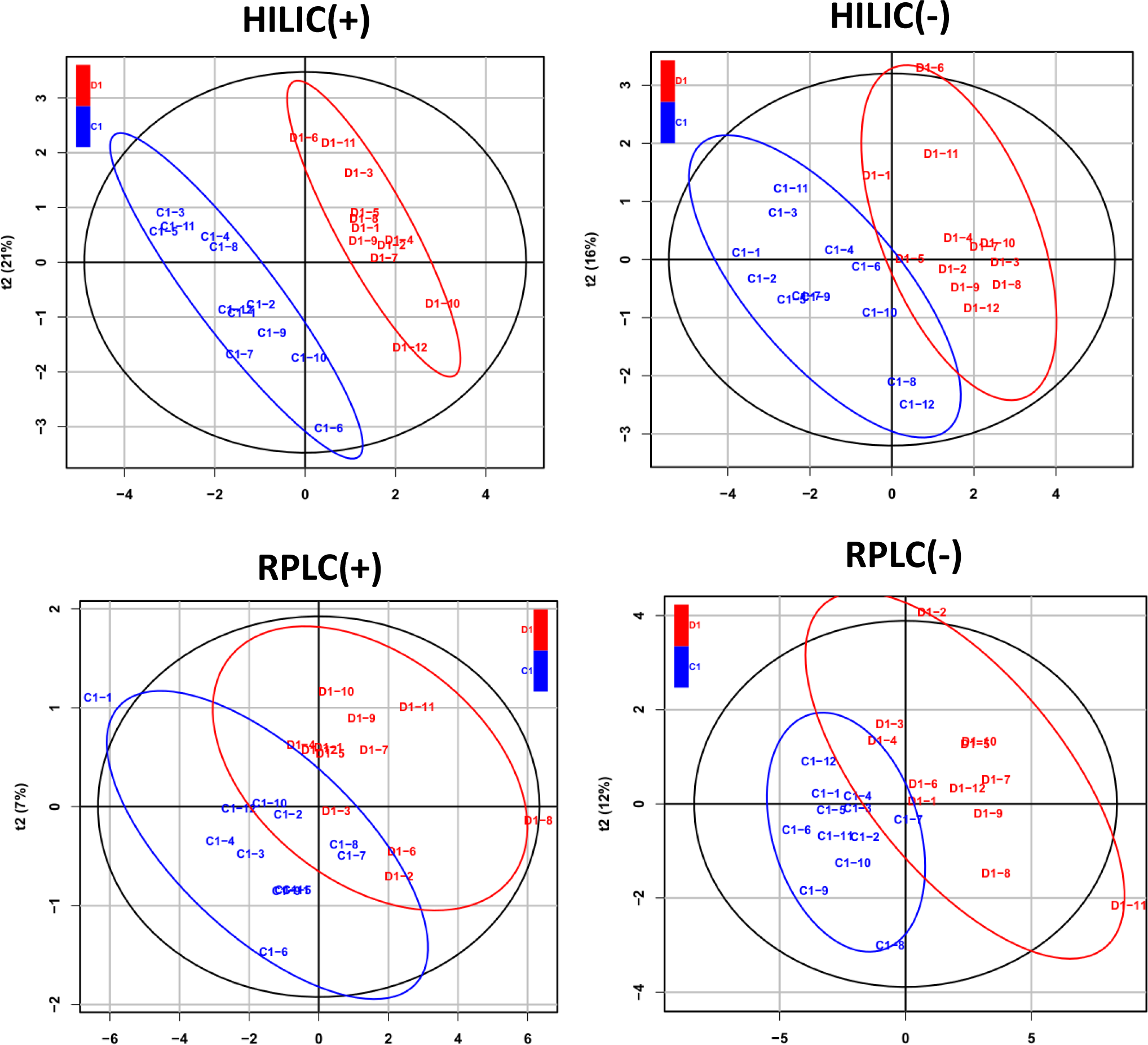
PLS-DA score scatter plots of the datasets presenting discrimination of female organisms during reproductive stage. Each dot represents a gammarid organism sample at a specific reproductive stage (C1 in blue and D1 in red). Score plots show significant separation between the different reproductive stage and among the different multi-omics data (p<0.05). The ellipse for each group represented hotelling’s T-squared 95% confidence interval.

### 3.5. Lipid structure refinement using EAD

To go deeper into structural determination of the molecules identified as features in positive ion mode, Electron-Activated Dissociation (EAD) was used. Among the 16 most relevant features in the RPLC(+) dataset, only 5 compounds were confidently identified through CID MS/MS database matching, while the rest remain unknown. Interestingly, out of those 6 lipids, the phospholipid PC 36:3 was identified and highlighted as an important feature in both RPLC(+) and RPLC(-) datasets. This lipid was identified at the same RT in positive and negative ion mode, and the obtained trends are identical, increasing the confidence in the results (Fig. 8). In CID, the prominent and intense fragment observed is the loss of a phosphocholine fragment at *m/z* 184.07, which is a common fragment (polar head group) shared between phosphatidylcholines (PC) and sphingomyelins (SM). In negative ion mode, fragmentation allows the identification of fatty acids on the glycerol backbone. However, in the case of PC 36:3, several combinations of fatty acids produced the same identification score due to low fragment intensities, preventing a definitive identification. Hence, the information obtained through CID is limited to the total number of carbons and the number of unsaturation, lacking details about the specific combination of fatty acids, their sn-isomerism and the positions of unsaturation. The same limitation applies to other identified lipids, such as PC O-34:3, PC O-36:5, PC O-36:1, and TG 45:0. EAD was investigated and applied in a Zeno-MRM^HR^ method in RPLC(+) to either complete or elucidate the identification of those 16 relevant features. The EAD MS/MS spectrum of PC 36:3 exhibited two diagnostic fragment ions at *m/z* 224.10 containing a carbon atom and *m/z* 226.08 with an oxygen atom specific to the glycerol backbone (Fig. 9). This ion pair serves as a unique identifier for phosphatidylcholines and has been already reported in previous studies EAD fragmentation spectrum of the m/z 550.5092 precursor ion in RPLC(+)EAD fragmentation spectrum of the m/z 550.5092 precursor ion in RPLC(+) [28,29,31]. In the EAD MS/MS spectra, two fragment ions were observed at *m/z* 489.3 and 504.3. These ions correspond to the radical fragmentation of C18:1 on the glycerol’s carbon (*m/z* 489.3) and the neutral loss of the C18:2 fatty acid chain (*m/z* 504.3) at the sn-1 and sn-2 positions, respectively. Further exploration of the fatty acid chain lengths and double bond positions was conducted. EAD played a crucial role by providing essential information about the number of carbons within each chain and the position of the double bonds, as illustrated in Figure 10. Upon careful examination of ions within the *m/z* range of 500-780, a series of chain fragments (albeit of low intensity) originating from the intact precursor ion at *m/z* 784.58 was observed, showing sequential radical fragmentation loss of one carbon at a time from the fatty acid backbone. We anticipate three types of carbon losses: methyl radical loss (CH3, 15 Da) from the end chain carbon of fatty acids, methylene radical loss (CH2, 14 Da) indicating a simple covalent bond between carbons, and methylidyne radical loss (CH, 13 Da) revealing the presence of double bonds on the chain. These fragment ions, such as *m/z* 755.54 and *m/z* 741.52 indicated the presence of double bonds at the C9 and C12 positions of the fatty acid chain moieties. The identified fragments spanned the entire length of the fatty acid chains and confirmed the presence of oleic acid (FA 18:1, △9; CH_3_(CH_2_)_7_CH=CH(CH_2_)_7_CO-) and linoleic acid (FA 18:2, △9,12; CH_3_(CH_2_)_4_–CH=CH–CH_2_–CH=CH–(CH_2_)_7_–CO-) esterified to the glycerol backbone of PC. Additionally, the FA composition of PC was validated by the presence of the ions at *m/z* 281.5 and *m/z* 279.2 in negative CID MS/MS spectra. Consequently, the lipid species PC 36:3 (PC 18:1/18:2) was successfully identified as PC 18:1(9)/18:2(9,12) representing the main molecular species regarding other low intensity coeluting isobaric compounds (Fig. 5). This result aligned perfectly with our previous study [37]. Notably, the fatty acids in gammarids are predominantly omega-6 and in *trans* configuration [37].

**Figure 5:**
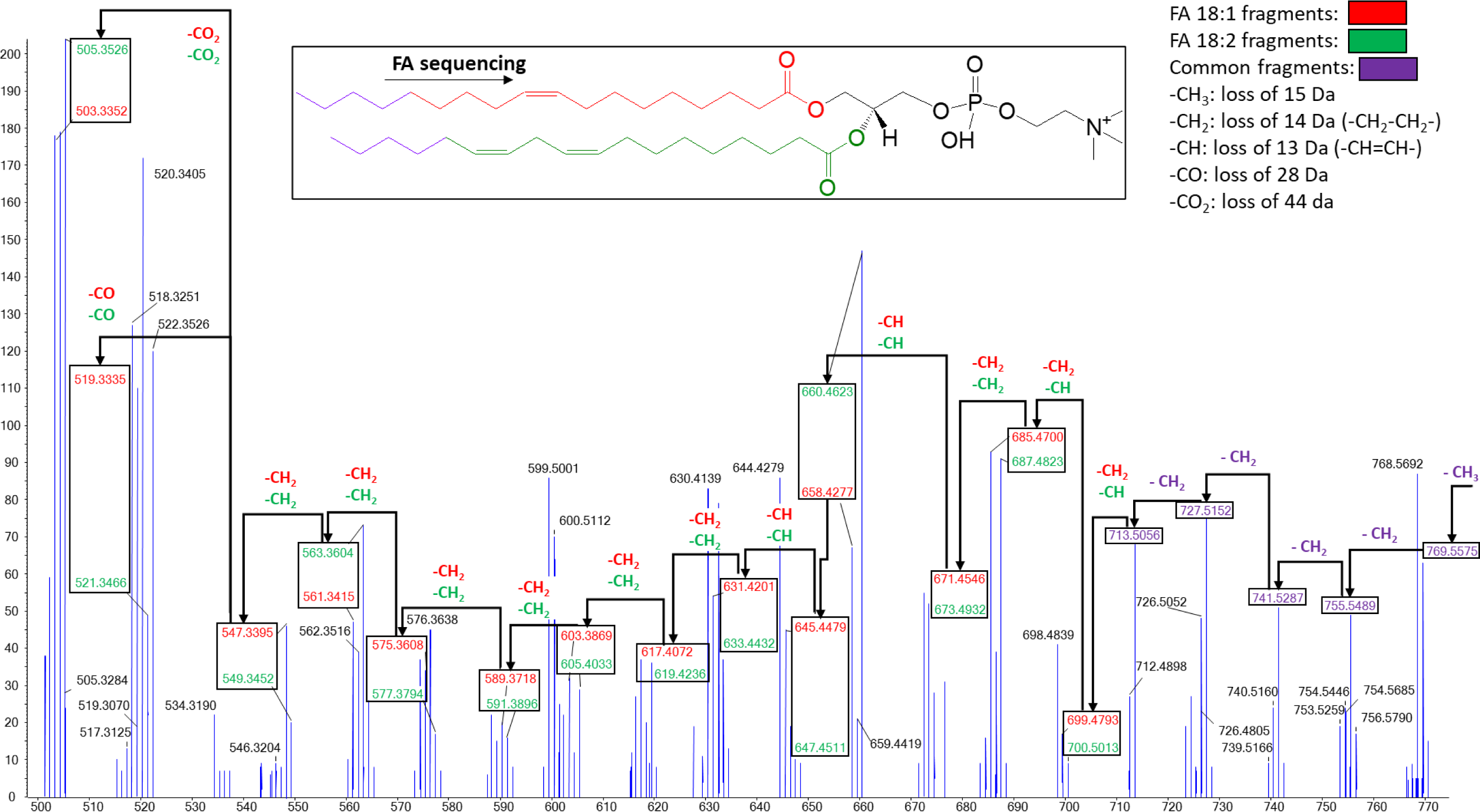
EAD MS/MS spectra of the discriminant feature PC 36:3, [M+H]^+^ *m/z* 784.58 from *m/z* range 500-770. Product ion spectra was recorded in positive ion mode with an electron kinetic energy of 15 eV. Black arrows indicate successive carbon losses in the fatty acid chains. The resulting fragments are color-coded as follows: red for C18:1, green for C18:2, and purple for fragments common to both fatty acids. The corresponding carbon loss is specified above each arrow using the same color code. Carbon losses can be -CH_3_ (15 Da), -CH_2_-(14 Da), -CH-(13 Da), -CO-(28 Da), or -CO_2_ (44 Da).

**Figure 6:**
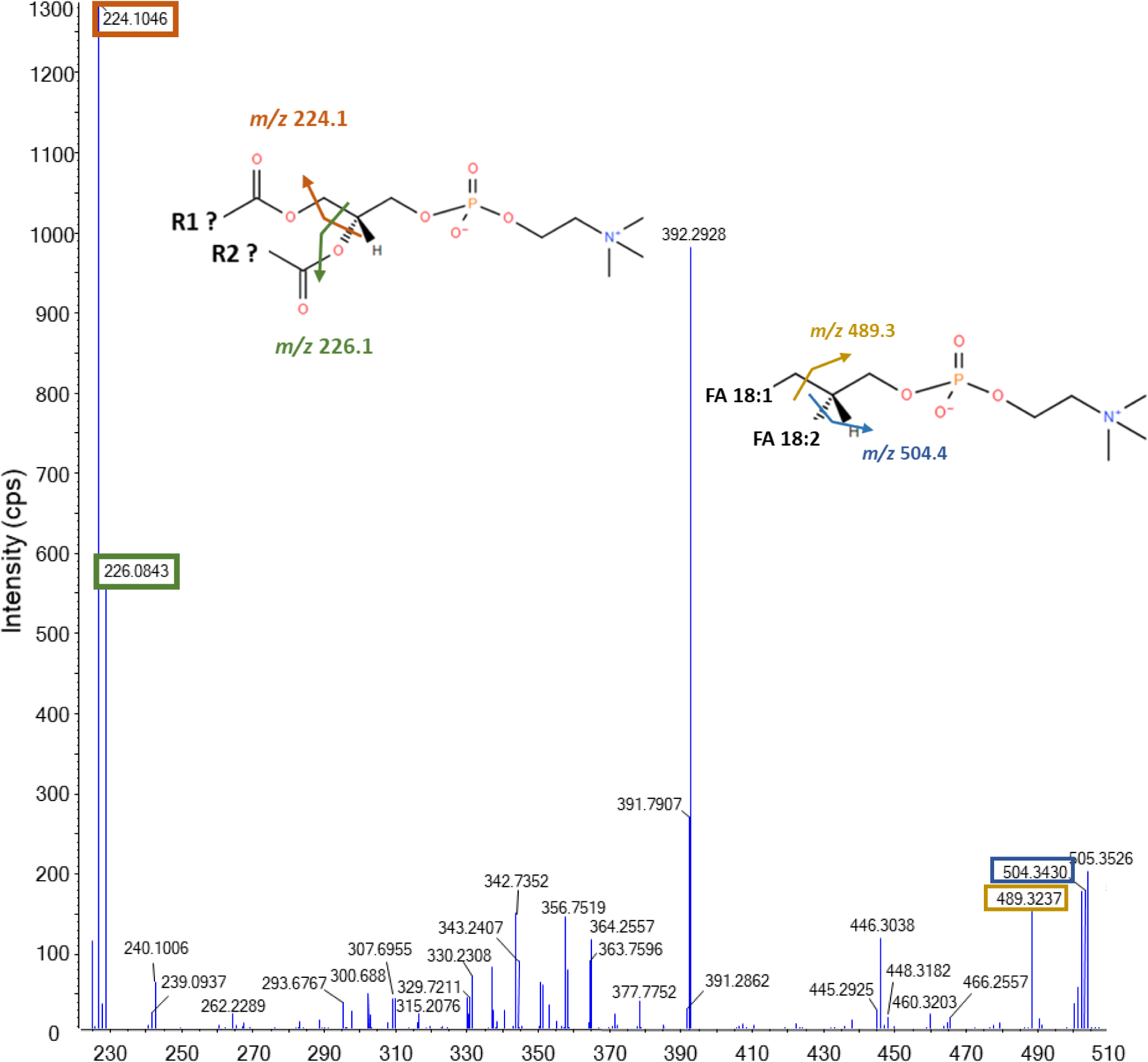
EAD MS/MS spectrum of PC 36:3 between 200-510 *m/z*. EAD generated several diagnostic fragment ions such as *m/z* 224.1046 (surrounded in orange) that contains a carbon atom and *m/z* 226.0843 (surrounded in green) that contains an oxygen atom for the glycerol backbone. This pair of ions enable the identification of phospholipid category. The ion *m/z* 429.3237 corresponds to the fragment ions formed at the cleavage site of the glycerol backbone leading to the loss of C18:1 and *m/z* 504.3430 for the loss of C18:2.

Concerning the other relevant features, those partially identified lipids were further confirmed and elucidated using EAD. Specifically, PC O-34:3 was identified as PC O-16:0/18:3(9,12,15), PC O-36:5 as PC O-16:0/20:5(5,8,11,14,17), PC O-36:1 as PC O-18:0/18:1(9), and TG 45:0 as TG 14:0/16:0/15:0. However, the unknown features that did not match any CID MS/MS spectra in MS-DIAL remained unidentified even after MS/MS EAD spectral database, that can be attributed to the limited comprehensiveness of the database.

### 3.6. Metabolite structure verification and confirmation by EAD

To highlight the benefits of EAD in the context of structural elucidation applied to metabolomics, we focused on the case of the CID MS/MS matched N-acetylglucosamine (considering all optical isomers). N-acetylglucosamine was matched through CID MS/MS database search to its [M + Na]^+^ adduct through four characteristic fragments: three neutral losses *m/z* 226.0604, 124.0365, 143.0301 and one radical fragmentation *m/z* 112.0500 illustrated in Figure 7’s top blue spectrum, alongside their respective fragmentation patterns. Despite achieving a decent matching score (879 MS-DIAL v.4 score), some ambiguities remained regarding the functional group on the hexose carbon n°2, purportedly identified as an acetamide. Since no CID fragmentation occurred on the carbon n°2 group, the structural confirmation of this group remained elusive, relying solely on the congruence of the global precursor ion mass and the match of the hexose fragments. This raises the possibility of the compound being an isobaric form of N-acetylglucosamine, distinguished exclusively by the carbon n°2 group. Consequently, our knowledge is limited to the molecular formula of the carbon n°2 group, C_2_H_4_NO, giving rise to various potential structures, as depicted in Figure 7, including 1-aminoethen-1-ol, acetimidic acid, or variations involving the linkage of acetamide to either the terminal carbon or the amide’s nitrogen. Additionally, the CID fragments of this adduct are relatively low in intensity, with the two fragments *m/z* 226.0604 and 143.0301 exhibiting relatively high ppm difference relative to the theoretical *m/z* value (Table 2). This discrepancy can be attributed to the impact of low fragments, leading to compromised mass accuracy. In general, the fragmentation of sodium adducts may be less efficient than that of proton adducts for several reasons. Sodium ions are heavier than proton ions, and as a result, the energy required to induce fragmentation may differ. Additionally, the chemical nature of the bond between sodium and the molecule can influence how easily fragmentation occurs. It is also possible that the fragmentation of sodium adducts leads to less stable ions, which could explain lower peak intensities in the mass spectrum. Given all of this information, the conclusion provided by CID are limited and should be approached with care in this case.

**Figure 7:**
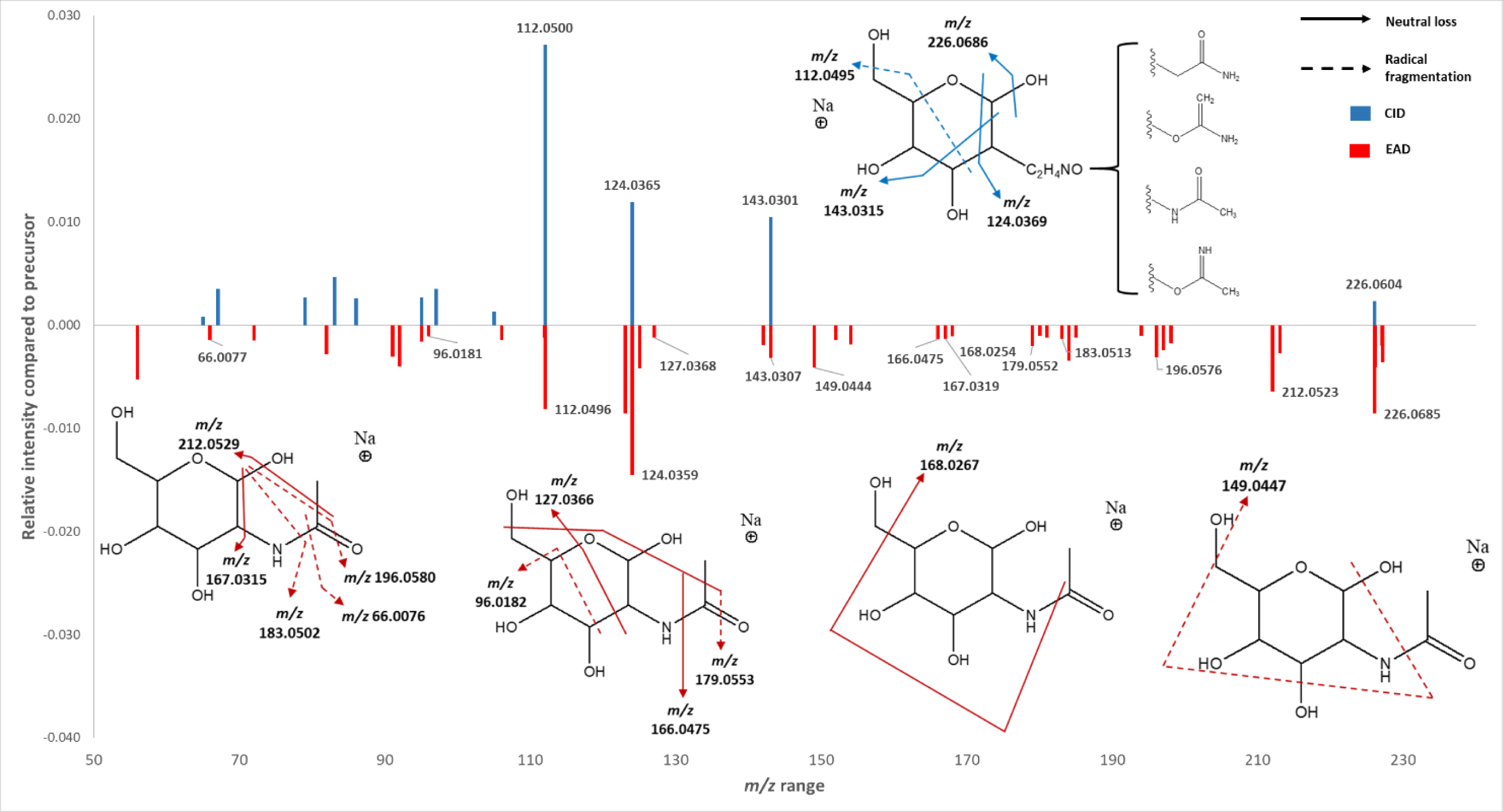
CID and EAD spectra comparison for putatively identified n-acetylglucosamine. Despite the absence of structural information on the supposed acetylamine group, the CID spectrum was matched with n-acetylglucosamine. The uncertainty arises from the group formula C_2_H_4_NO, which could correspond to multiple structural arrangements of the atoms. However, in the EAD spectrum, the ambiguity is resolved by the presence of additional characteristic fragments on this substructure. These fragments (*m/z* 212.0523, *m/z* 196.0576, *m/z* 183.0513, *m/z* 184.0344, *m/z* 184.0344, *m/z* 167.0319) confirm the identification as an acetylamine and, consequently, establish the global identification of n-acetylglucosamine (considering all optical isomers). The upper spectrum (in blue) represents the CID spectrum, with labeled fragments explained by fragmentation patterns using blue arrows. The lower spectrum (in red) corresponds to the EAD spectrum, and the labeled fragments are clarified through fragmentation patterns using red arrows. The solid arrows indicate neutral loss fragmentation, and the dotted arrows signify radical fragmentation. To enhance readability, the complex EAD fragmentation patterns are presented in four separate figures due to the abundance of fragments.

**Table 2.**
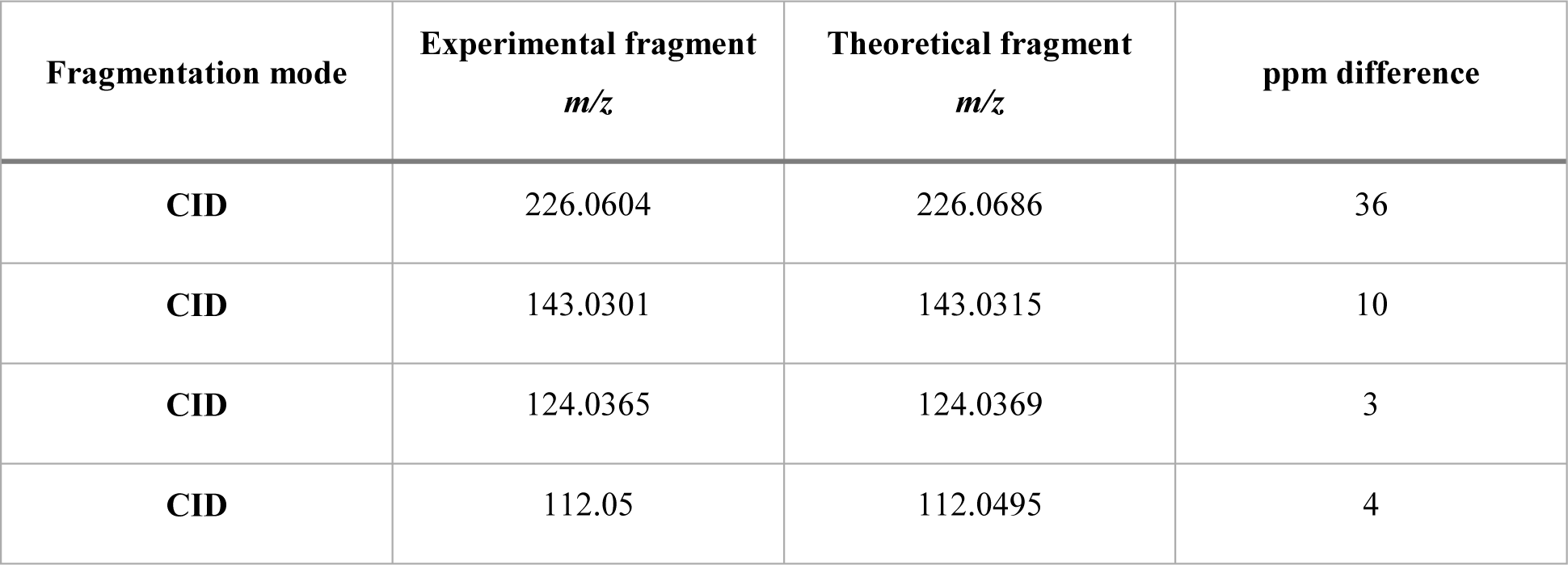
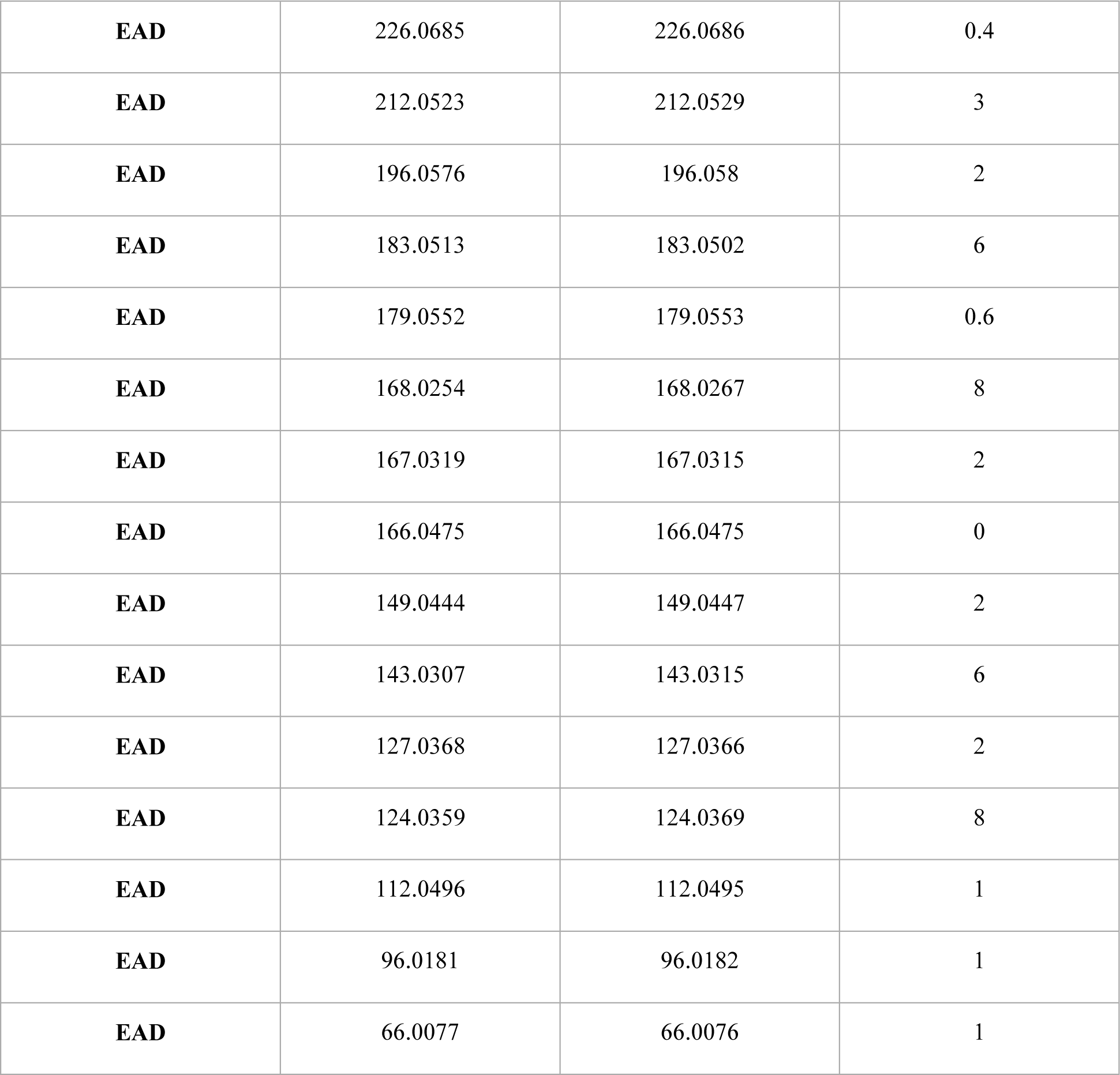
Fragment annotation of n-acetylglucosamine using CID and EAD. Each fragment annotated in Figure 7 for n-acetylglucosamine are referenced with their experimental and theoretical *m/z* values. A ppm difference is calculated for each one.

| Fragmentation mode | Experimental fragment<br>$m/z$ | Theoretical fragment<br>$m/z$ | ppm difference |
| --- | --- | --- | --- |
| **CID** | 226.0604 | 226.0686 | 36 |
| **CID** | 143.0301 | 143.0315 | 10 |
| **CID** | 124.0365 | 124.0369 | 3 |
| **CID** | 112.05 | 112.0495 | 4 |
| **EAD** | 226.0685 | 226.0686 | 0.4 |
| **EAD** | 212.0523 | 212.0529 | 3 |
| **EAD** | 196.0576 | 196.058 | 2 |
| **EAD** | 183.0513 | 183.0502 | 6 |
| **EAD** | 179.0552 | 179.0553 | 0.6 |
| **EAD** | 168.0254 | 168.0267 | 8 |
| **EAD** | 167.0319 | 167.0315 | 2 |
| **EAD** | 166.0475 | 166.0475 | 0 |
| **EAD** | 149.0444 | 149.0447 | 2 |
| **EAD** | 143.0307 | 143.0315 | 6 |
| **EAD** | 127.0368 | 127.0366 | 2 |
| **EAD** | 124.0359 | 124.0369 | 8 |
| **EAD** | 112.0496 | 112.0495 | 1 |
| **EAD** | 96.0181 | 96.0182 | 1 |
| **EAD** | 66.0077 | 66.0076 | 1 |

When applying EAD to the same molecule, additional fragments were generated, particularly in the high *m/z* range between 150 and 220, as illustrated in the bottom EAD red spectrum of Figure 7. Despite having lower intensity than CID fragments, the EAD fragments provided more detailed structural information, especially concerning the carbon n°2 group while still encompassing the CID fragments. Eleven fragments were manually identified and annotated in Figure 7, with a deviation of less than 10 ppm from the theoretical values (Table 2). Among them, five fragments (*m/z* 212.0523, 196.0576, 183.0513, 167.0319, 66.0077), predominantly associated with the carbon n°2 group, played a crucial role in confirming the identification of acetamide and, consequently, establishing the overall identification of N-acetylglucosamine. The presence of a terminal carbon on the carbon n°2 group was confirmed by the neutral loss leading to the *m/z* 212.0523 fragment, while radical fragmentations leading to *m/z* 196.0576, 183.0513, and 66.0077 confirmed the presence of a carbonyl group (Figure 7). Additionally, the neutral loss resulting in *m/z* 167.0319 confirmed the presence of nitrogen directly linked to the hexose carbon n°2 (Figure 7). Several other informative fragments, induced by both radical and non-radical fragmentation processes, further supported the global identification of N-acetylglucosamine. Fragments at *m/z* 179.0552, 168.0254, 166.0475, 149.0444, 127.0368, and 96.0181, originating partly from hexose fragmentation, not only verified the correct identification of the hexose portion but also contributed additional information about the carbon n°2 group to augment the previous findings (*m/z* 179.0552, 168.0254, 166.0475, 149.0444) (Figure 7).

### 3.7. Database-free approach by SIRIUS for structure investigation using EAD

To go further into the structural elucidation of the metabolites and lipids features in gammarids we combined EAD and an *in-silico* free database approach with SIRIUS [21–27]. Currently, most of the databases used for the structural elucidation of metabolites are based on CID MS/MS spectra. Since EAD fragmentation is more recent, only MS-DIAL version 5.1 software integrates EAD for lipid identification. Nevertheless, this database remains partial, and potential identifications are limited to ‘known unknowns,’ meaning the compounds in the database. However, database-free solutions that provide structural propositions through the investigation or prediction of fragmentation spectra are an engaging alternative. Two strategies are considered: either MS/MS spectra from known compounds are algorithmically predicted and tentatively matched to experimental spectra, or the experimental spectra are computationally investigated to reconstruct the molecular structure [45]. The first approach relies on ‘known unknowns,’ which lacks completeness compared to MS/MS spectral databases. On the other hand, the latter approach relies on deep-learning methods such as support vector machines, enabling the modelling between fragmentation spectra and molecular fingerprints after large-scale model training. Databases can subsequently be searched after structure prediction to find already existing compounds. If the prediction is missing from databases, valuable information on substructures and chemical classes is still obtained. This approach was chosen as the full metabolome of *G. fossarum* is not documented yet, and thus no information is available regarding the biomolecular content.

For compound identification, we used MS/MS database from CSI:FingerID available in SIRIUS [26]. This *in-silico* approach relies on the combination of the precursor exact mass, the fragmentation spectrum and eventually the RT to predict molecular formulas using the exact mass and the isotopic pattern, and structure propositions using MS/MS spectraNext, SIRIUS was employed on the 10 most discriminant features from HILIC(+) and the 16 from RPLC(+) using CID and EAD spectra to provide structural insights. In HILIC(+), we confirmed the identification of N-acetylglucosamine (considering all optical isomers) for the [M + Na]^+^ adduct at *m/z* 244.07. Among the 9 HILIC(+) unknown features initially referred to as retention time_ *m/z* of precursor ion, 5 were confidently and consistently predicted with Tanimoto scores above 75% and CSI:FingerID scores above −70 (Table S5): 15.7_202.07, 10.3_204.09, 14.8_434.16, 6.4_461.28, 13.9_573.32. Relevant insights into the remaining 4 unknown structures were obtained. Two of them were likely small oligopeptides (14_442.27 and 14.8_549.27), an another one a hydroxy acid (5.3_217.07), and the last one might be an oligopeptide with several serine (12.4_690.32). Regarding RPLC(+), the previously putatively identified lipids were confirmed by SIRIUS with strong Tanimoto and CSI:FingerID scores: PC O-34:3 (95%, −17), PC O-36:5 (99%, −9.1), PC O-36:1 (91%, −48), TG 45:0 (100%, −0.4), and PC 36:3 (95%, −11) (Table S5). Four of the unknown features received highly reliable structure predictions with Tanimoto scores above 93% and CSI:FingerID above −20: 5_284.29 as an amine with a possible alkyne (95%, −19), 5.4_565.4 as alloxanthin (93%, −21), 10.6_880.76 as TG 53:7 (98%, −9.2), and 9.4_969.74 as TG 58:8 (97%, −13). Due to the occurrence of certain alkynes in plants, their potential presence in gammarids could be elucidated. Furthermore, Alloxanthin exhibits functions associated with alkynes. However, TG 58:8 retention time is inconsistent with the number of carbons and unsaturation, indicating incomplete structure prediction but still valuable. Six unknowns were provided hints about substructures rather than relevant structures: 9.7_548.5, 9.7_550.51, 9.4_587.55, 5.4_605.4, 9.5_698.26, 10.2_750.41. One feature did not yield a relevant prediction, proposing only oligopeptides, which are inconsistent with the retention time. Table S5 contains this information, including the proposed structure name, adduct, and interpretation/comments about substructures.

EAD-based structure prediction for HILIC(+) confirmed the annotation of N-acetylglucosamine at both *m/z* 244.07 for the [M + Na]^+^ adduct and *m/z* 204.08 for the [M – H_2_O + H]^+^ adduct (Table 8). This dual confirmation is logical as both features share the same RT and trends between stages. While the [M-H_2_O + H]^+^ N-acetylglucosamine MS/MS spectrum didn’t yield a relevant structure prediction in CID, EAD provided a clearer identity through a more informative MS/MS spectrum (Table S6). N-acetylglucosamine is the monomer of chitin, the primary component of arthropods’ exoskeletons, making its presence consistent due to the reproductive cycle’s modification during moulting [46]. Another intriguing discovery was the structurally similar feature at *m/z* 434.1649 in HILIC(+), suggesting the addition of a hexose to N-acetylglucosamine that underwent ortho hydroxy group etherification. This compound formation mirrors the chitin synthesis pathway, where phosphorylated N-acetylglucosamine molecules are de-phosphorylated and assembled to form chitin [46]. However, as this pathway has not yet been demonstrated in *G. fossarum*, further studies are required. In HILIC(+), six features were predicted with the same structure as CID, with scores not significantly altered. The SIRIUS algorithm penalizes fragmentations like EI fragmentation in ESI, affecting the overall scores. Since EAD fragments are generally less intense than CID fragments, their contribution to the score is minor. Half of the significant features in HILIC(+) are probably oligopeptides, while the other half comprise sugars and hexose-type molecules.

In RPLC(+), the five previously MS/MS matched features retained the correct structure, supporting the putative identifications: PC O-34:3 (96%, −18), PC O-36:5 (91%, −31), PC O-36:1 (96%, −38), TG 45:0 (87%, −72), and PC 36:3 (99%, −9.5) (Table S6). Among the initially unknowns, four obtained the same structure prediction as CID: Alloxanthin (93%, −29), TG 53:7 (96%, 6.4%), TG 58:8 (97%, −13), and a phosphonic acid-like compound (48%, −82) (Table S6). However, caution is needed with the phosphonic acid prediction due to imperfect scores. Interestingly, EAD suggested alloxanthin as the plausible structure for the *m/z* 605.39 feature, contradicting the CID prediction. This feature appears to be a [M + H_2_O + Na]^+^ adduct, explaining its identical RT and trend among groups as the [M + H]^+^ alloxanthin adduct at 565.40 m/z. Alloxanthin is a carotenoid acting as a protector molecule against photooxidation and can be biologically produced or absorbed through plant and algae consumption [47,48].

In conclusion, EAD provides additional information for the interpretation of fragmentation spectra compared to CID. This fragmentation mode can validate or invalidate the annotations established through machine learning methodologies. All the EAD fragmentation spectra both in HILIC(+) and RPLC(+) are available in supplementary data (Figure S11-36).

## Conclusion

In conclusion, SWATH-DIA has demonstrated significant benefits for characterizing the metabolome and lipidome of the crustacean amphipod *Gammarus fossarum* during the reproductive stage, in comparison of IDA/DDA acquisition mode. The Zeno pulsing effect, facilitated by the Zeno trap, enhances detection sensitivity, leading to a more comprehensive molecular fingerprint crucial for biologically relevant features. These benefits are evidenced by a minimum 5-fold increase in fragment intensity for over half of the fragments in all datasets (52%). Moreover, 78% of features exhibit at least twice as many fragments per fragmentation spectrum with Zeno pulsing. These results advocate for employing Zeno pulsing alongside the SWATH-DIA approach, which synergistically combines extensive coverage and enhanced detection sensitivity. Additionally, these advancements are underscored by a substantial 57% increase in the number of confident putative identifications based on MS/MS databases across all datasets, emphasizing the methodology’s significant contribution to improving compound identification accuracy and reliability.

Regarding compound annotation, the use of EAD by the targeted approach ZENO-MRM^HR^ has significantly contributed to elucidating specific lipid species in contrast of CID MS/MS spectra. For instance, PC 36:3, previously identifiable only through CID in positive ion mode, was successfully elucidated as PC 18:1(9)/18:2(9,12) using EAD in one single acquisition (one polarity). However, EAD-based identifications rely on the MS/MS reference database in MS-DIAL 5.1, and this database-driven approach has limitations as it restricts identifications to “known unknowns” in the database, introducing bias into the identification process. To address this limitation, a database-free approach employing SIRIUS has been explored. Through a Multiple Kernel Learning approach for structure prediction, previously unknown features in both HILIC(+) and RPLC(+) datasets have provided valuable insights into potential structures and substructures. Despite these advancements, caution is warranted when interpreting results from machine learning tools since annotation has to be confirmed by synthetic standards.

### CRediT authorship contribution statement

**Thomas Alexandre Brunet**: Conceptualization, Methodology, Formal analysis, Investigation, Writing – original draft, Visualization. **Yohann Clément**: Conceptualization, Software, Formal Analysis, Supervision. **Valentina Calabrese**: Investigation, Supervision. **Jérôme Lemoine**: Resources. **Olivier Geffard**: Investigation, Resources. **Arnaud Chaumot**: Investigation, Resources. **Davide Degli-Esposti**: Investigation, Resources. **Arnaud Salvador**: Resources, Writing – review & editing. **Sophie Ayciriex**: Conceptualization, Project administration, Funding acquisition, Writing – review & editing.

## Acknowledgments

The authors thank the French Ministry of Higher Education, Research, and Innovation (Ministère de l’Enseignement Supérieur, de la Recherche et de l’Innovation) for the PhD fellowship of Thomas Alexandre Brunet. Valentina Calabrese was supported by a post-doctoral fellowship of the SENS research funding of the Université Claude Bernard Lyon 1. This work was supported by the French National Agency Research (Young investigator Grant ANR-18-CE34-0008) and performed within the framework of the EUR H2O’Lyon (ANR-17-EURE-0018) of Université de Lyon (UdL), within the program “Investissements d’Avenir”. The authors also benefitted from the French GDR “Aquatic Ecotoxicology” framework which aims at fostering stimulating scientific discussions and collaborations for more integrative approaches.

**Appendix A. Supplementary data**

