## Supplemental figures and Tables for "Concomitant investigation of crustacean amphipods lipidome and metabolome during molting stage by Zeno SWATH Data-Independent Acquisition coupled with Electron Activated Dissociation and machine learning"

**Appendix A - Supplemental data**

Figure S1: SWATH variable windows assays for the four LC-MS/MS methods. *m/z* density histograms : normalized density of the precursor ions (blue line) according to the window's width (orange). (A) RPLC in positive ionization mode, (B) RPLC in negative ionization mode, (C) HILIC in positive ionization mode and (D) HILIC in negative ionization mode.

Figure S2. MS/MS spectra comparison of the putatively identified histamine when using the Zeno pulsing.

Figure S3. MS/MS spectra comparison of the putatively identified tyrosine when using the Zeno pulsing.

Figure S4. MS/MS spectra comparison of the putatively identified abrine when using the Zeno pulsing.

Figure S5. MS/MS spectra comparison of the putatively identified 6-oxopurine when using the Zeno pulsing.

Figure S6. MS/MS spectra comparison of the putatively identified TG 16:0/18:1/20:1 when using the Zeno pulsing.

Figure S7. MS/MS spectra comparison of the putatively identified LCP 18:2 when using the Zeno pulsing.

Figure S8. PCA scores of each dataset considering the samples (C1 in red and D1 in green), the QC (in cyan) and the blanks (in blue) before univariate test filtering. All datasets were log10 transformed and standardized.

Figure S9. Box plots of the putatively identified PC 36:3 in RPLC(+) and RPLC(-) comparing the log10 transformed intensity in C1 and D1. The boxes, error bars, medians and averages were calculated using the 12 values per stage. The C1 stage is represented in blue and the D1 in yellow. The difference was tested by a univariate test and a p-value below 0.0001 was observed hence the **** in both case.

Figure S10. EAD spectrum of putatively annotated PC 18:1/18:2 acquired in RPLC(+) mode between 50 and 790 m/z. Additionally, a magnified view, scaled up 10 times, focuses on the m/z range from 220 to 785.

Figure S11. EAD fragmentation spectrum of the m/z 202.0690 precursor ion in HILIC(+).

Figure S12. EAD fragmentation spectrum of the *m/z* 204.0875 precursor ion in HILIC(+).

Figure S13. EAD fragmentation spectrum of the *m/z* 217.0679 precursor ion in HILIC(+).

Figure S14. EAD fragmentation spectrum of the *m/z* 244.0793 precursor ion in HILIC(+).

Figure S15. EAD fragmentation spectrum of the *m/z* 434.1649 precursor ion in HILIC(+).

Figure S16. EAD fragmentation spectrum of the *m/z* 442.2668 precursor ion in HILIC(+).

Figure S17. EAD fragmentation spectrum of the *m/z* 461.2764 precursor ion in HILIC(+).

Figure S18. EAD fragmentation spectrum of the *m/z* 549.2684 precursor ion in HILIC(+).

Figure S19. EAD fragmentation spectrum of the *m/z* 573.3248 precursor ion in HILIC(+).

Figure S20. EAD fragmentation spectrum of the *m/z* 690.3202 precursor ion in HILIC(+).

Figure S21. EAD fragmentation spectrum of the *m/z* 284.2942 precursor ion in RPLC(+).

Figure S22. EAD fragmentation spectrum of the *m/z* 548.5032 precursor ion in RPLC(+).

Figure S23. EAD fragmentation spectrum of the *m/z* 550.5092 precursor ion in RPLC(+).

Figure S24. EAD fragmentation spectrum of the *m/z* 565.4033 precursor ion in RPLC(+).

Figure S25. EAD fragmentation spectrum of the *m/z* 587.5485 precursor ion in RPLC(+).

Figure S26. EAD fragmentation spectrum of the *m/z* 605.3956 precursor ion in RPLC(+).

Figure S27. EAD fragmentation spectrum of the *m/z* 679.4302 precursor ion in RPLC(+).

Figure S28. EAD fragmentation spectrum of the *m/z* 698.2554 precursor ion in RPLC(+).

Figure S29. EAD fragmentation spectrum of the *m/z* 742.5756 precursor ion in RPLC(+).

Figure S30. EAD fragmentation spectrum of the *m/z* 750.4069 precursor ion in RPLC(+).

Figure S31. EAD fragmentation spectrum of the *m/z* 766.5737 precursor ion in RPLC(+).

Figure S32. EAD fragmentation spectrum of the *m/z* 774.6373 precursor ion in RPLC(+).

Figure S33. EAD fragmentation spectrum of the *m/z* 782.7221 precursor ion in RPLC(+).

Figure S34. EAD fragmentation spectrum of the *m/z* 784.5848 precursor ion in RPLC(+).

Figure S35. EAD fragmentation spectrum of the *m/z* 880.7608 precursor ion in RPLC(+).

Figure S36. EAD fragmentation spectrum of the *m/z* 969.7362 precursor ion in RPLC(+).

**Supplementary tables**

Table S1: Zeno-MRM$^{HR}$-EAD parameters in HILIC(+) for features exhibiting a VIP above 1.5

Table S2: Zeno-MRM$^{HR}$-EAD parameters in RPLC(+) for features exhibiting a VIP above 1.5

Table S3: Filtering steps on the datasets.

Table S4: Summary of the PLS-DA models performance parameters.

Table S5: SIRIUS prediction under CID fragmentation.

Table S6: SIRIUS prediction under EAD fragmentation.

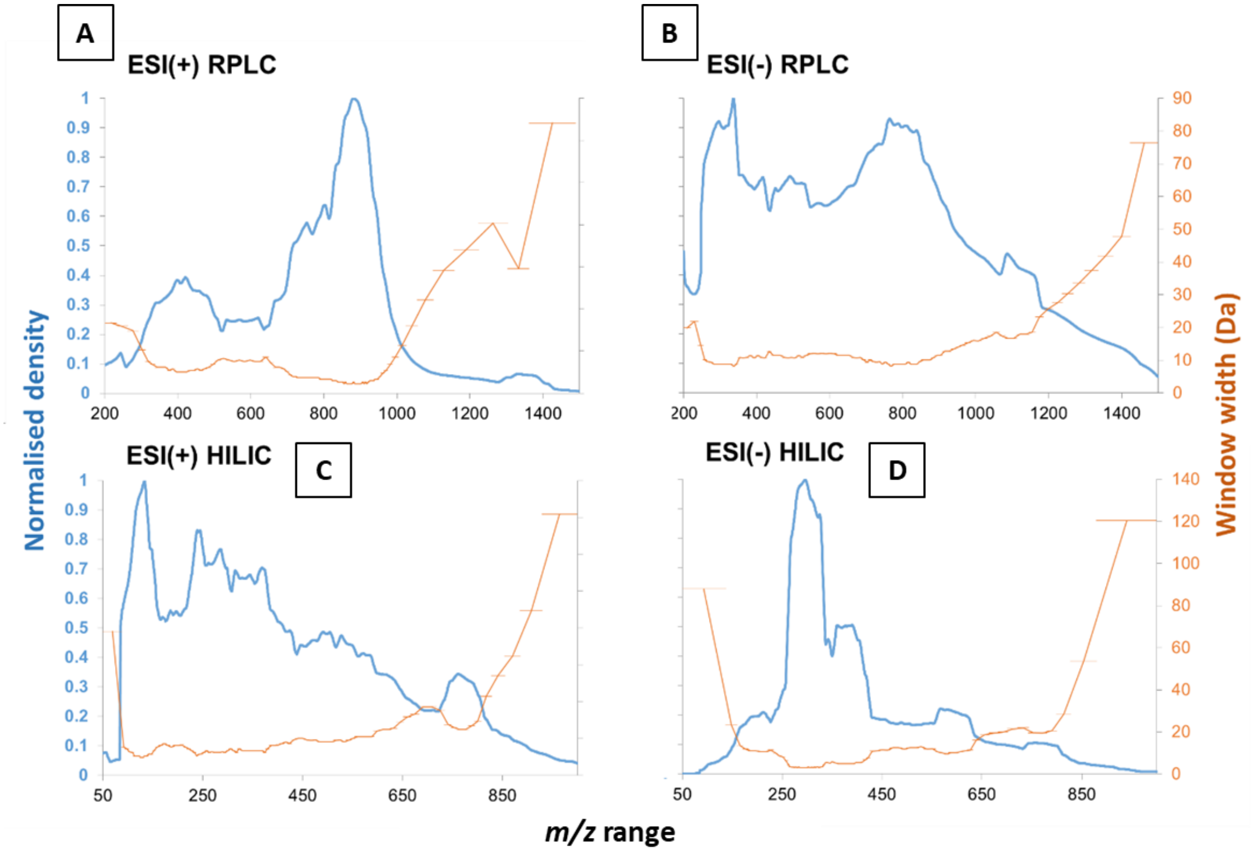
A
ESI(+) RPLC
B
ESI(-) RPLC
C
ESI(+) HILIC
D
ESI(-) HILIC
Normalised density
Window width (Da)
m/z range

Figure S1.

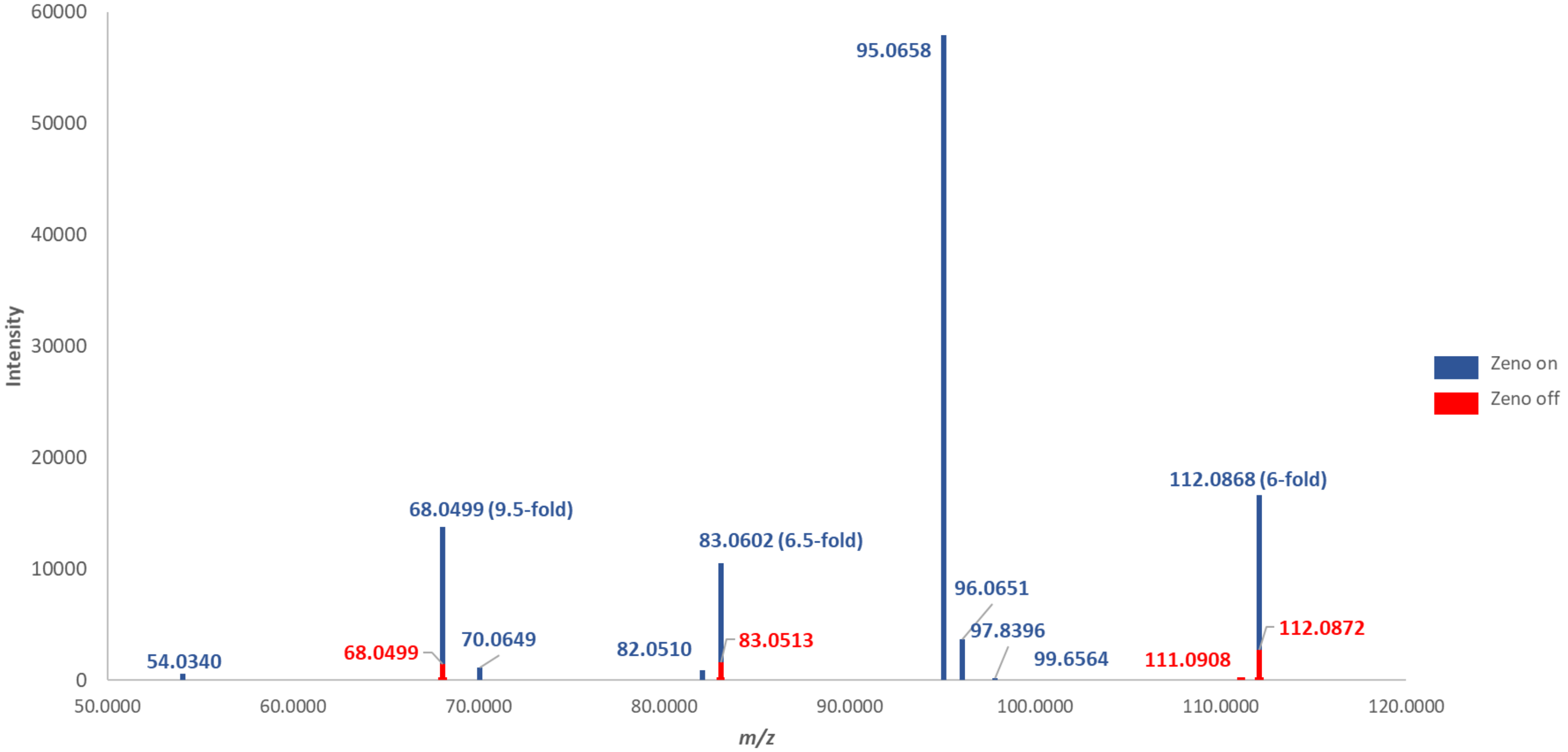
Intensity
60000
50000
40000
30000
20000
10000
0
95.0658
112.0868 (6-fold)
68.0499 (9.5-fold)
83.0602 (6.5-fold)
96.0651
97.8396
112.0872
70.0649
83.0513
68.0499
82.0510
54.0340
99.6564
111.0908
50.0000
60.0000
70.0000
80.0000
90.0000
100.0000
110.0000
120.0000
m/z
Zeno on
Zeno off

Figure S2.

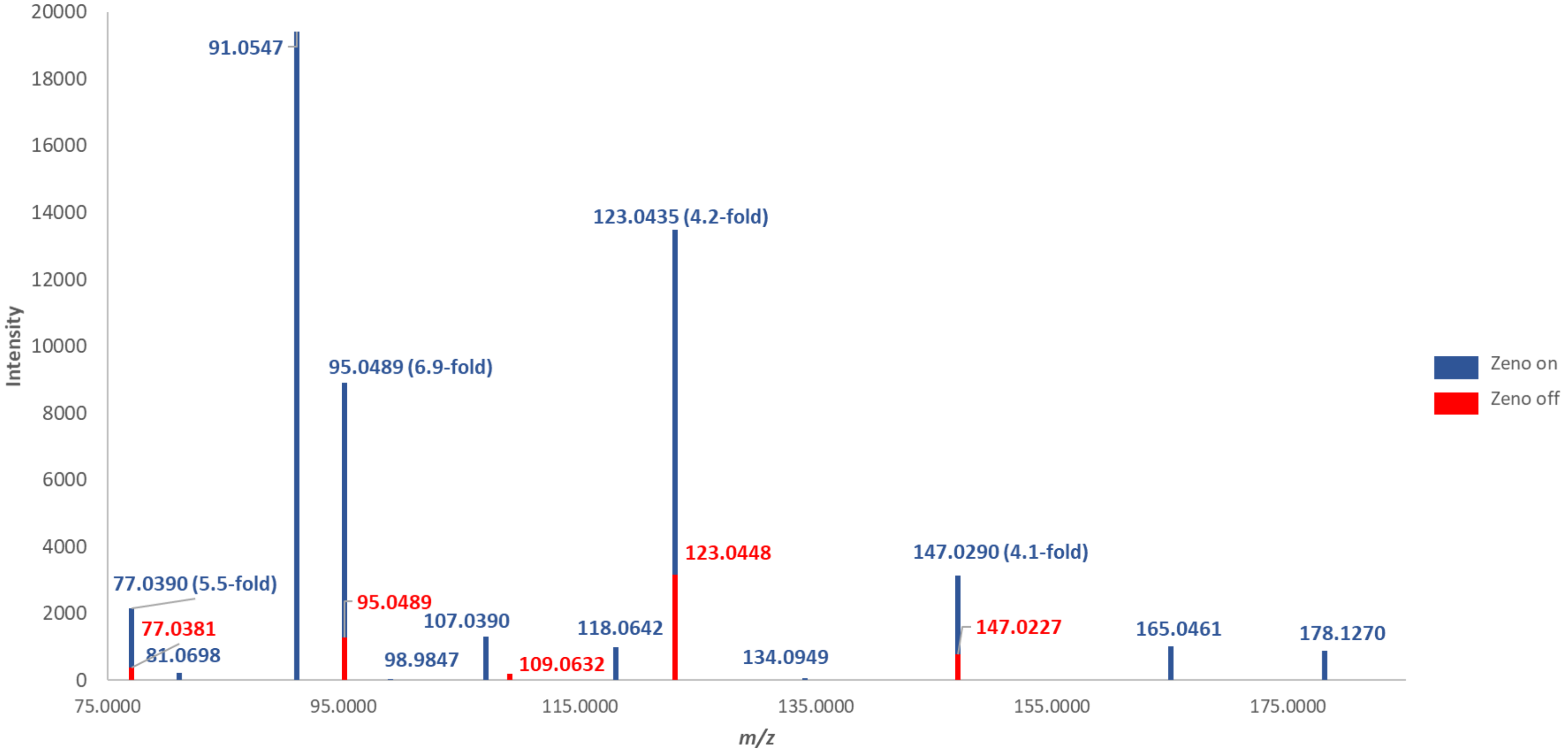

Intensity
20000
18000
16000
14000
12000
10000
8000
6000
4000
2000
0
91.0547
123.0435 (4.2-fold)
95.0489 (6.9-fold)
123.0448
147.0290 (4.1-fold)
77.0390 (5.5-fold)
95.0489
107.0390
118.0642
147.0227
165.0461
178.1270
77.0381
81.0698
98.9847
109.0632
134.0949
75.0000
95.0000
115.0000
135.0000
155.0000
175.0000
m/z
Zeno on
Zeno off


Figure S3.

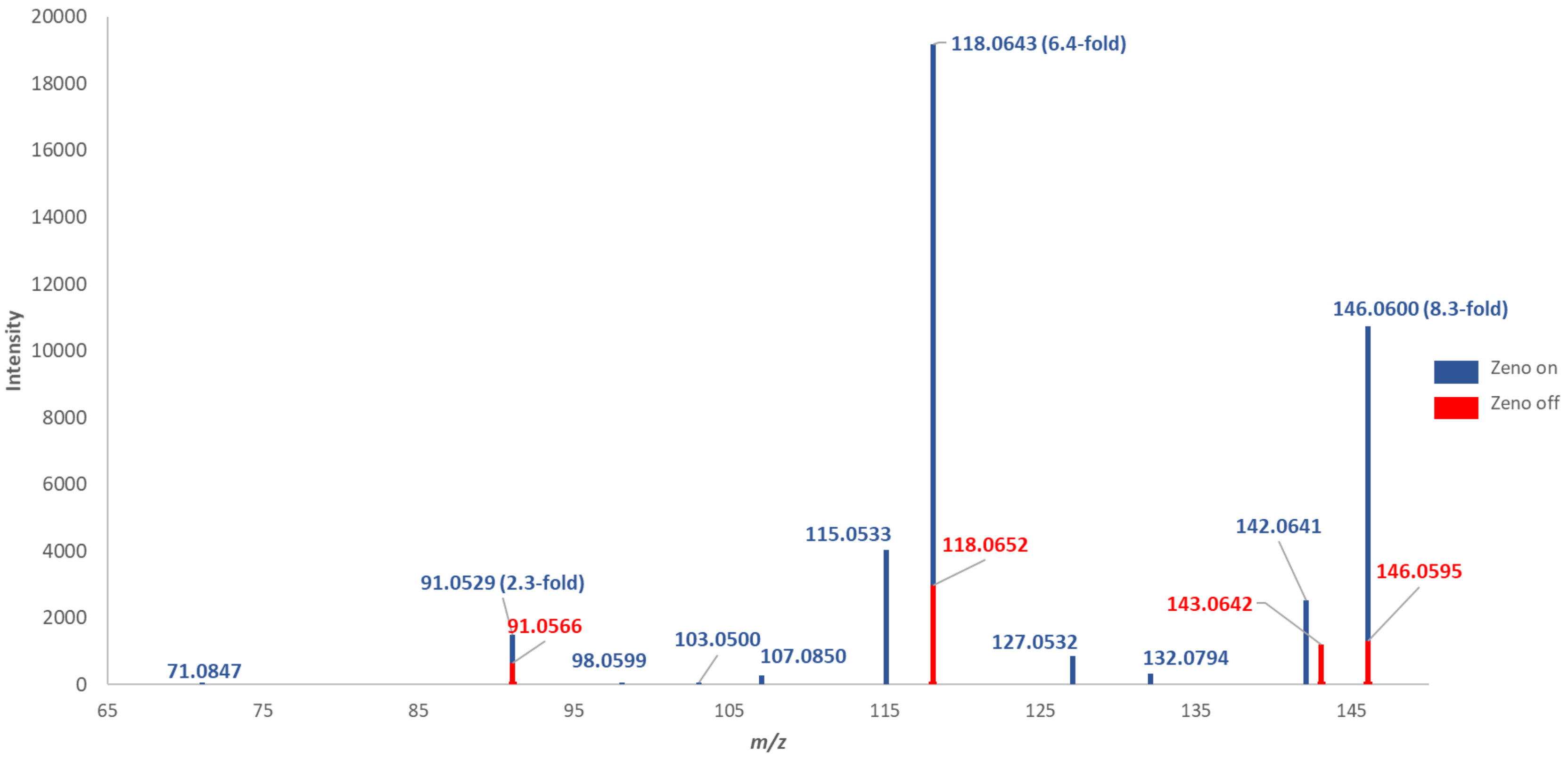

20000
18000
16000
14000
12000
10000
8000
6000
4000
2000
0
Intensity
118.0643 (6.4-fold)
146.0600 (8.3-fold)
Zeno on
Zeno off
115.0533
118.0652
142.0641
146.0595
91.0529 (2.3-fold)
143.0642
91.0566
103.0500
107.0850
127.0532
132.0794
98.0599
71.0847
65
75
85
95
105
115
125
135
145
m/z


Figure S4.

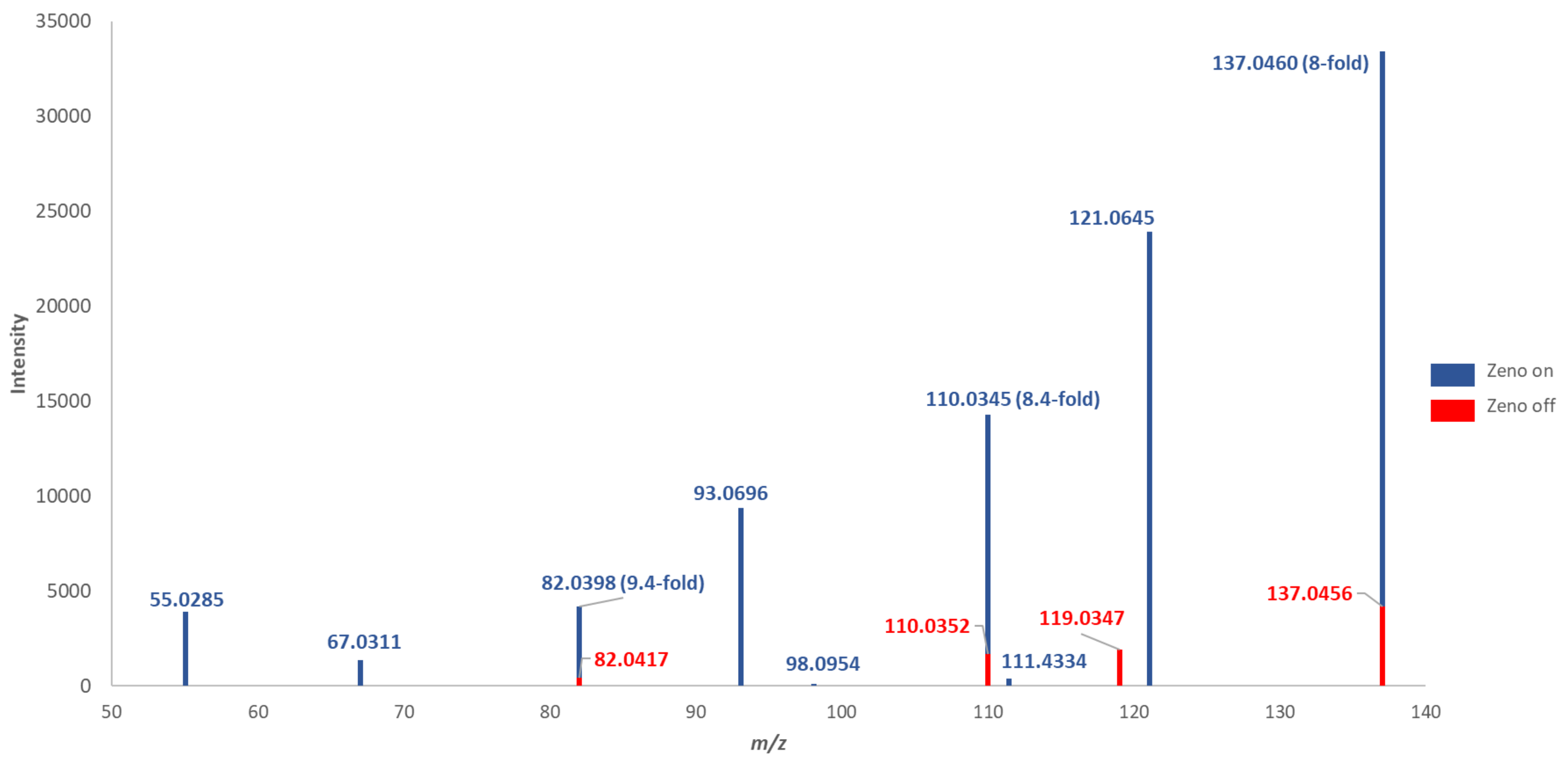

35000
30000
25000
20000
15000
10000
5000
0
Intensity
50
60
70
80
90
100
110
120
130
140
m/z
55.0285
67.0311
82.0398 (9.4-fold)
82.0417
93.0696
98.0954
110.0345 (8.4-fold)
110.0352
111.4334
119.0347
121.0645
137.0460 (8-fold)
137.0456
Zeno on
Zeno off


Figure S5.

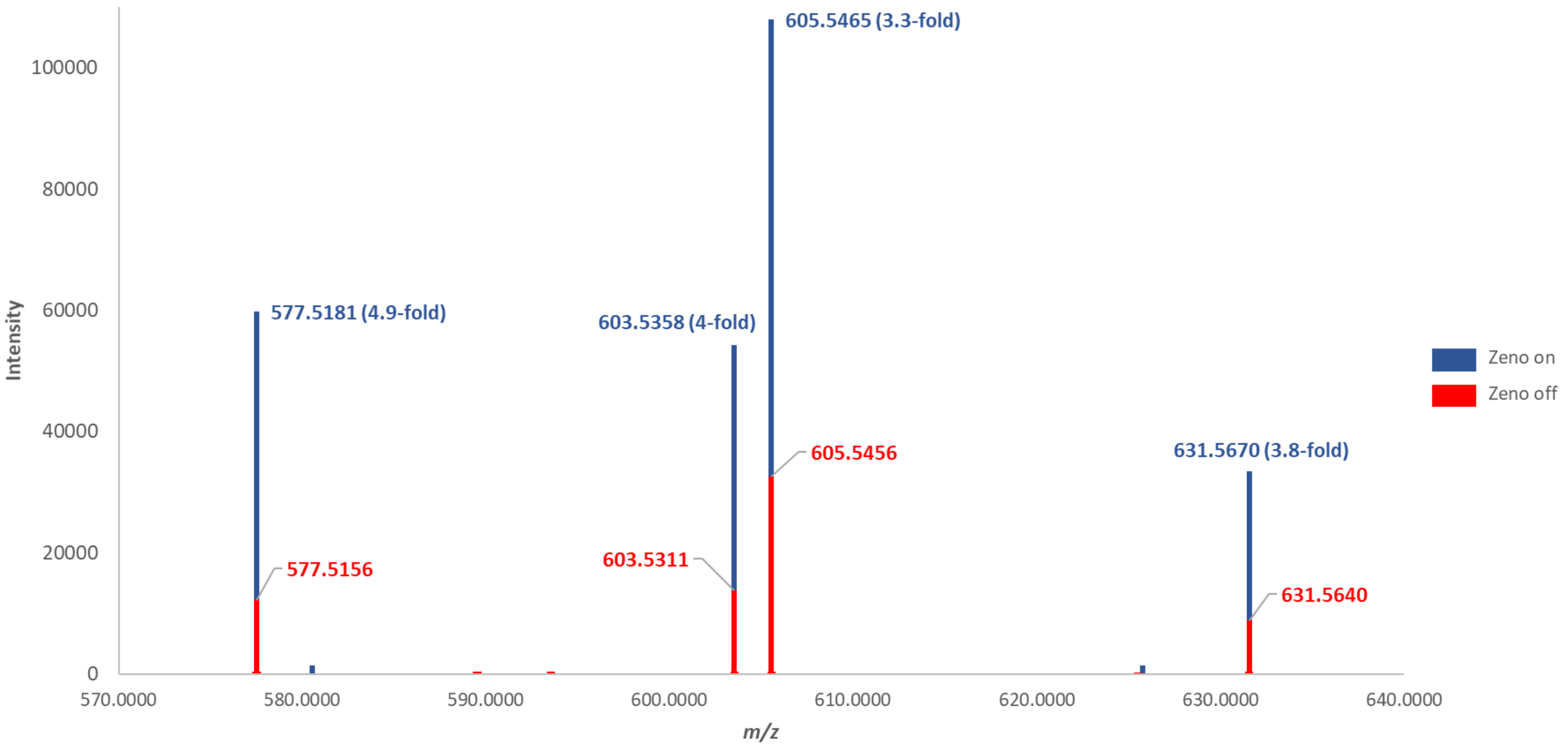
605.5465 (3.3-fold)
100000
80000
60000
577.5181 (4.9-fold)
603.5358 (4-fold)
Intensity
Zeno on
Zeno off
40000
605.5456
631.5670 (3.8-fold)
20000
577.5156
603.5311
631.5640
0
570.0000
580.0000
590.0000
600.0000
610.0000
620.0000
630.0000
640.0000
m/z

Figure S6.

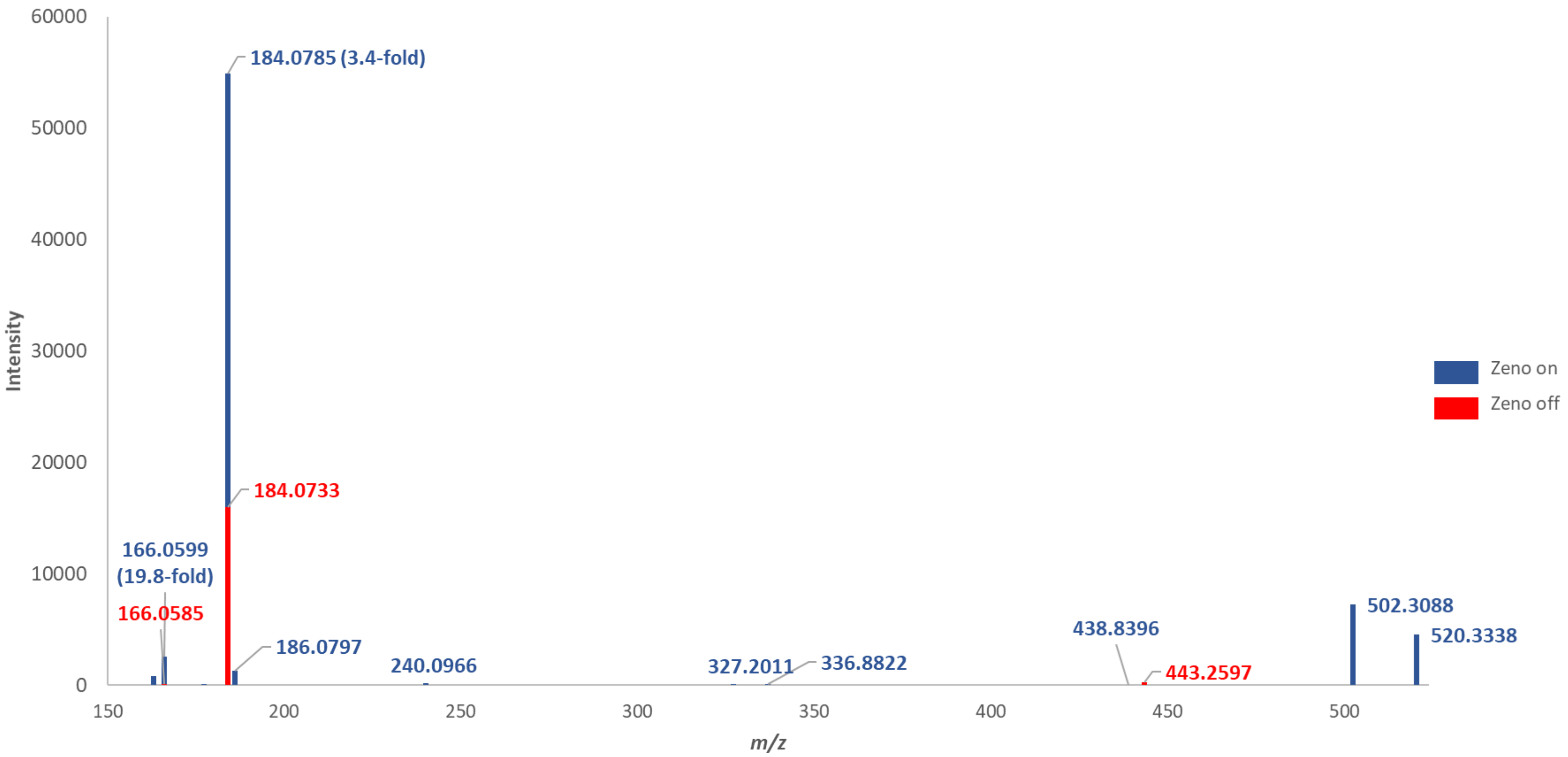
60000
50000
40000
30000
20000
10000
0
Intensity
184.0785 (3.4-fold)
184.0733
166.0599
(19.8-fold)
166.0585
186.0797
240.0966
327.2011
336.8822
438.8396
443.2597
502.3088
520.3338
Zeno on
Zeno off
150
200
250
300
350
400
450
500
m/z

Figure S7.

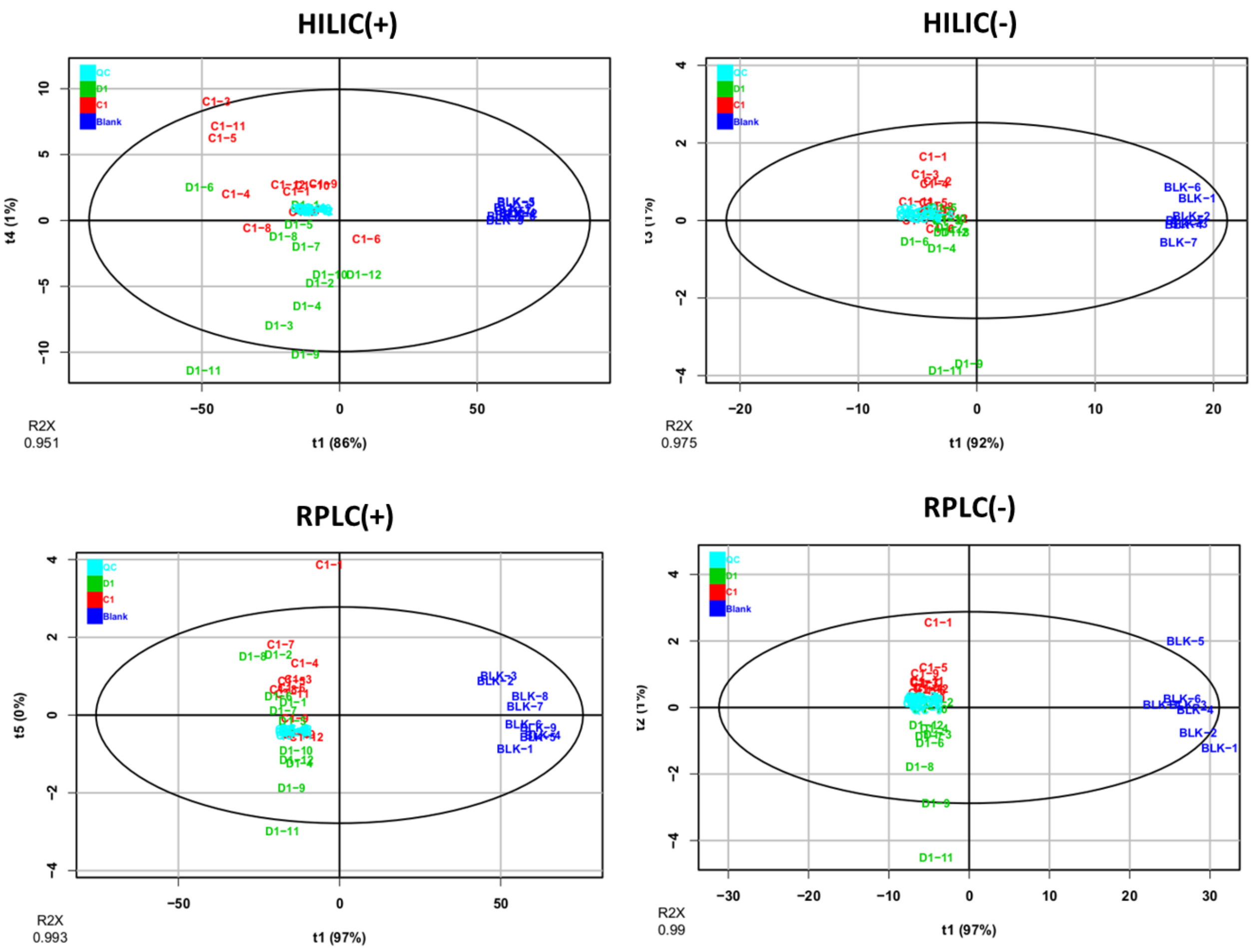
HILIC(+)
QC
D1
C1
Blank
t4 (1%)
t1 (86%)
R2X
0.951
HILIC(-)
t1 (92%)
R2X
0.975
RPLC(+)
t5 (0%)
t1 (97%)
R2X
0.993
RPLC(-)
t1 (97%)
R2X
0.99

Figure S8.

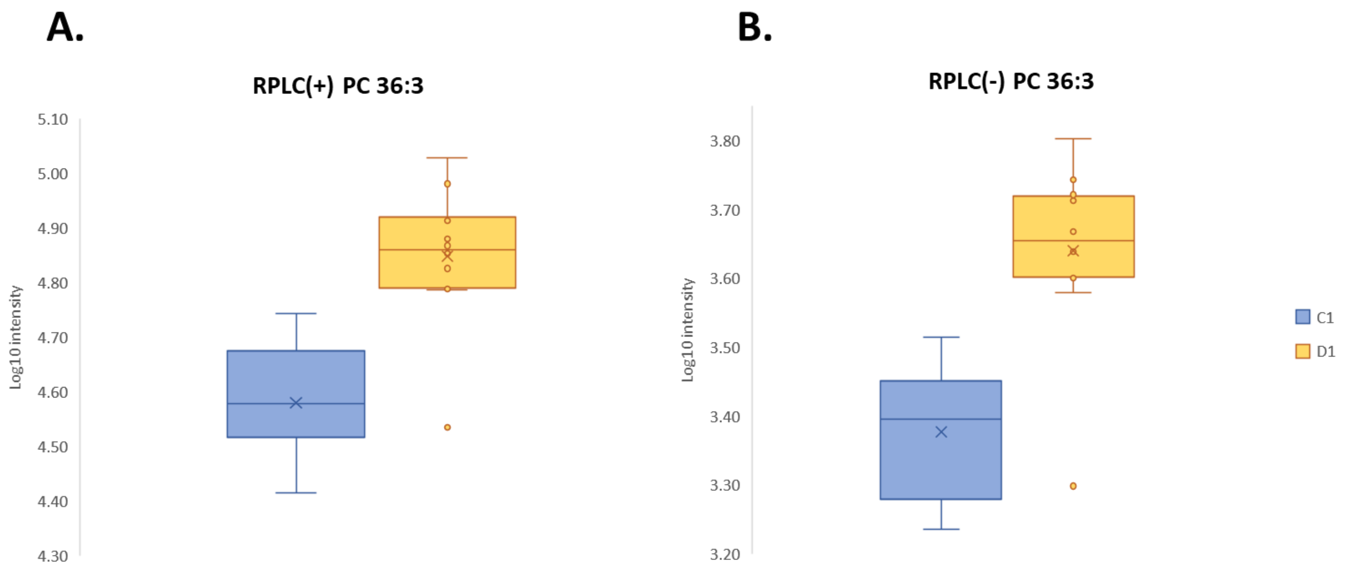

A.
RPLC(+) PC 36:3
Log10 intensity
5.10
5.00
4.90
4.80
4.70
4.60
4.50
4.40
4.30
B.
RPLC(-) PC 36:3
Log10 intensity
3.80
3.70
3.60
3.50
3.40
3.30
3.20
C1
D1


Figure S9.

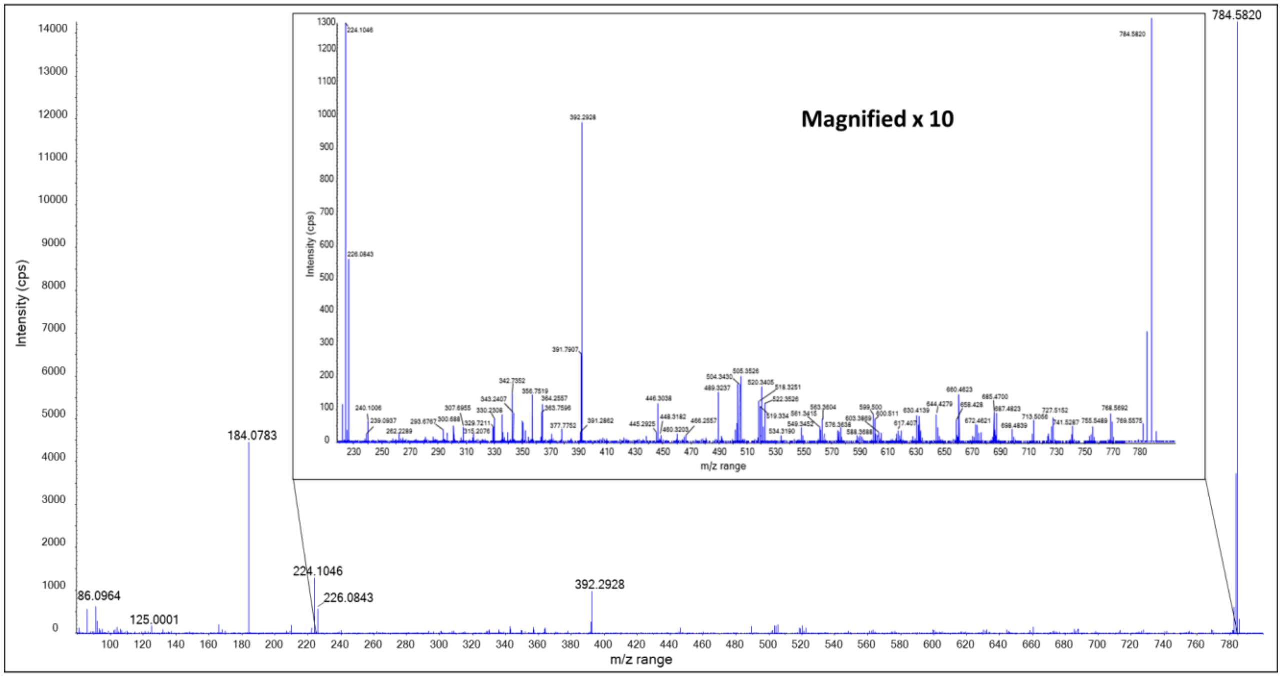


Figure S10.

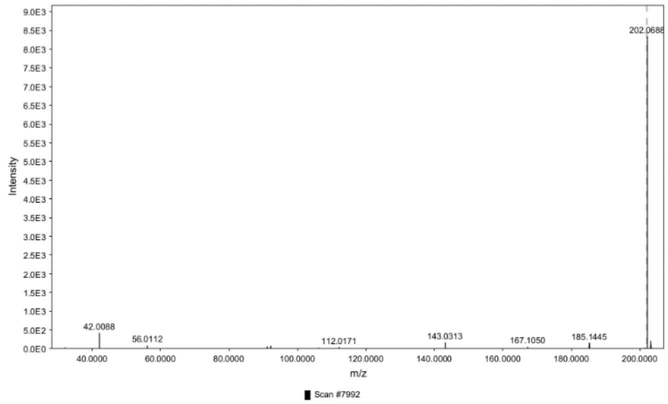


Figure S11

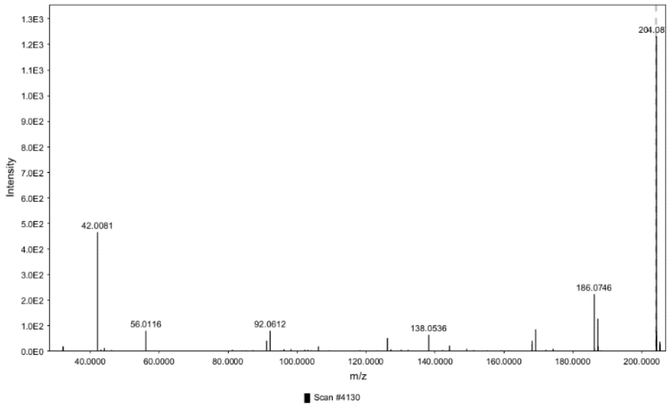


Figure S12

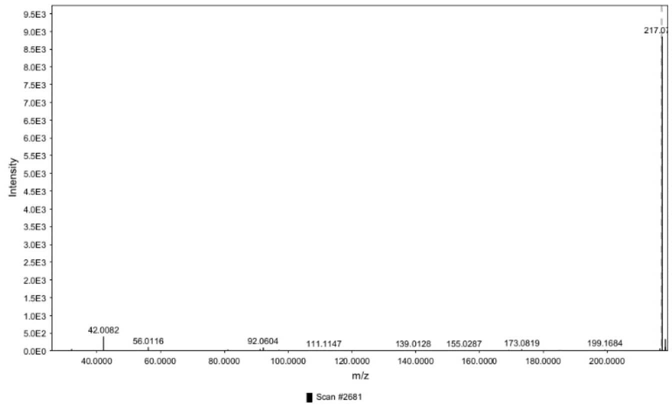


FigureS13

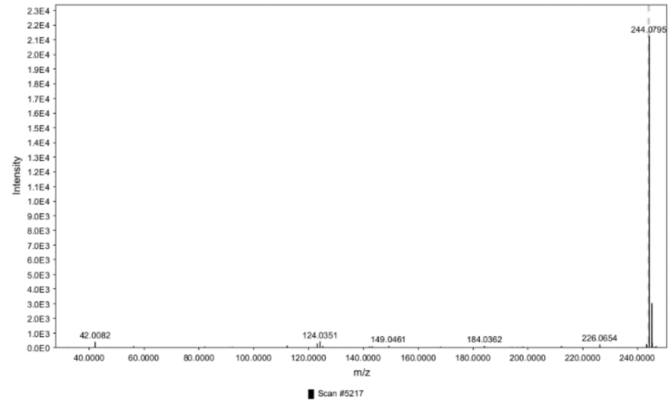


Figure S14

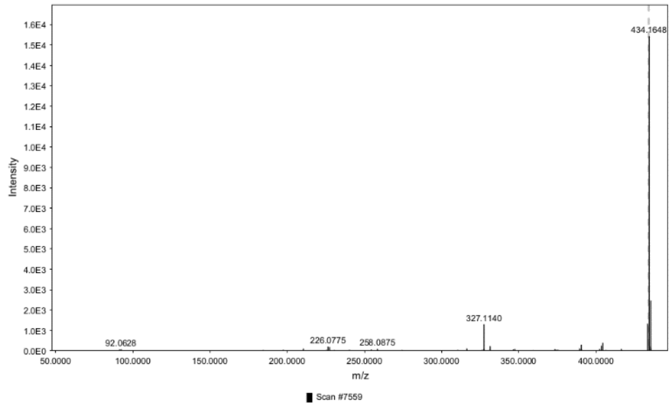


FigureS15

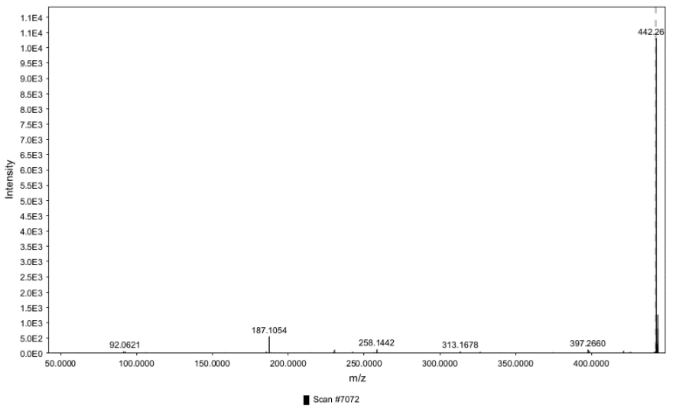


Figure S16

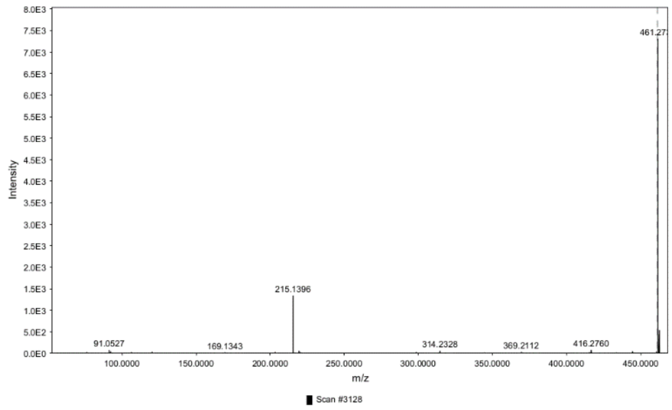


FigureS17

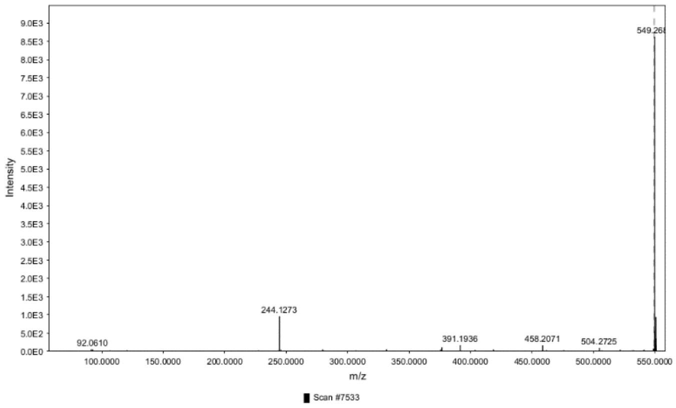


Figure S18

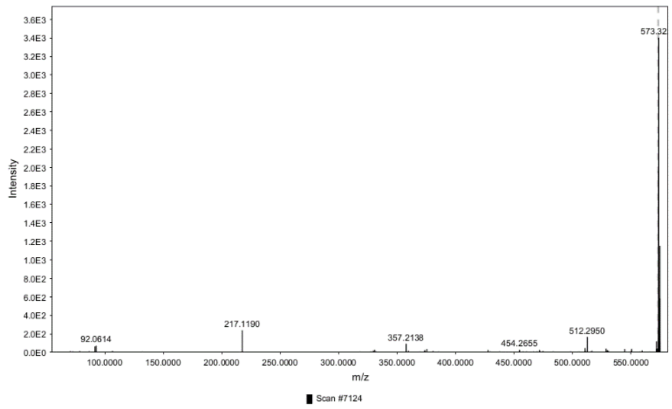


FigureS19

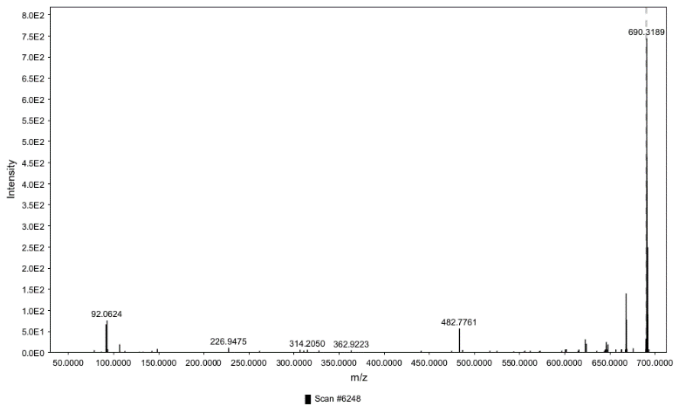


Figure S20

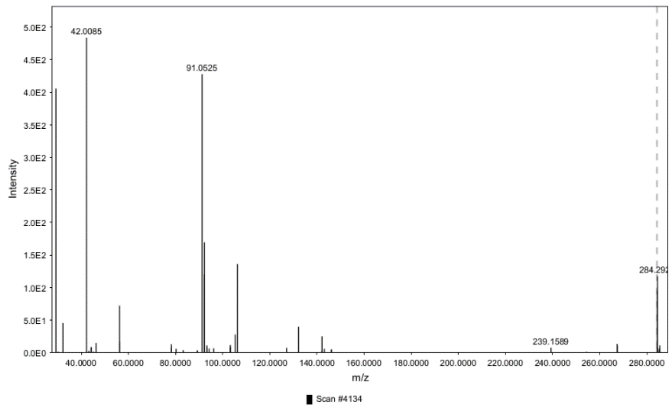


FigureS21

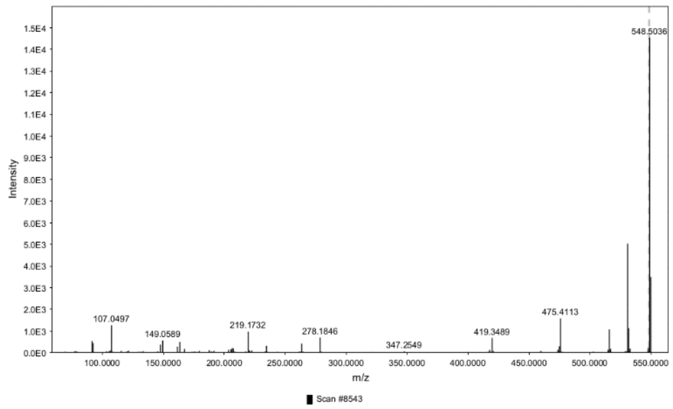


Figure S22

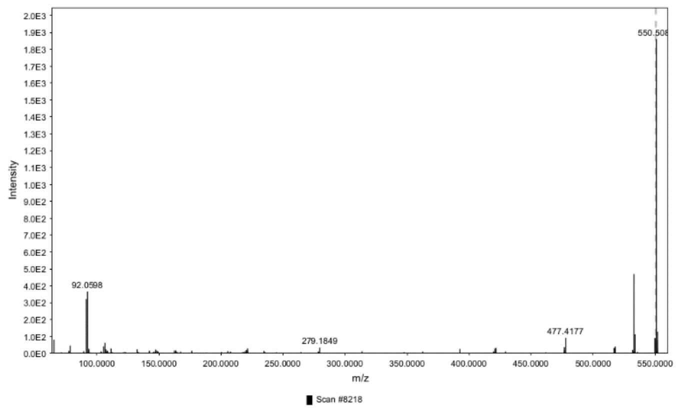


FigureS23

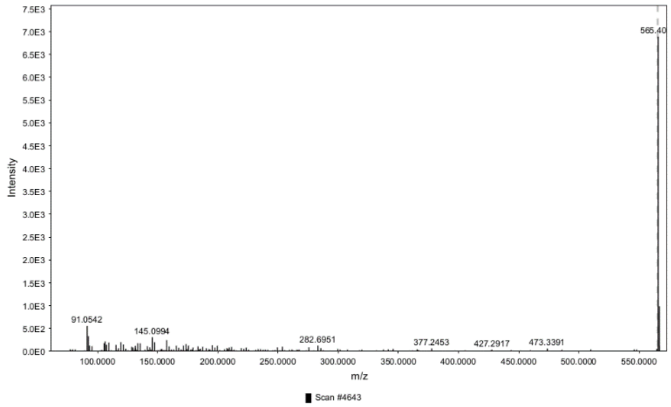


Figure S24

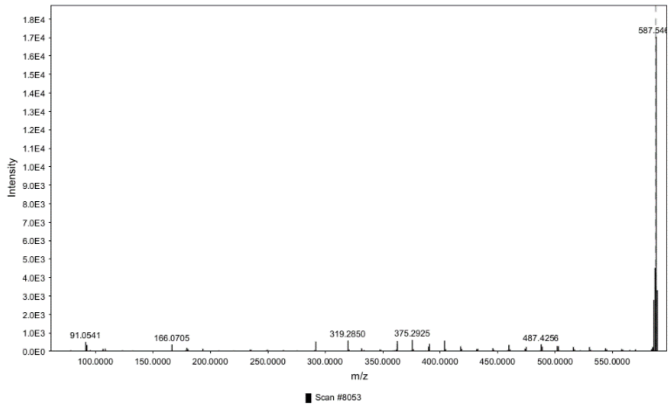


FigureS25

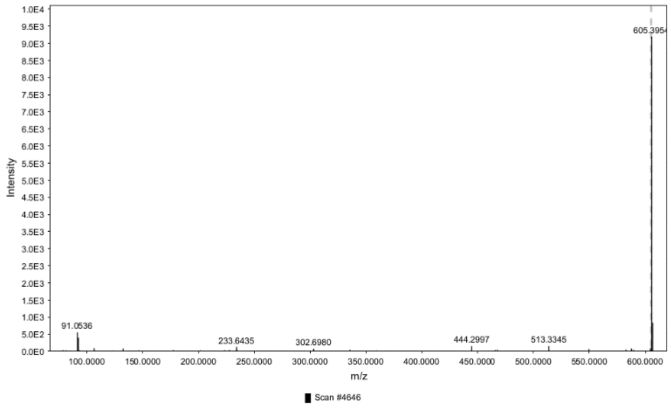


Figure S26

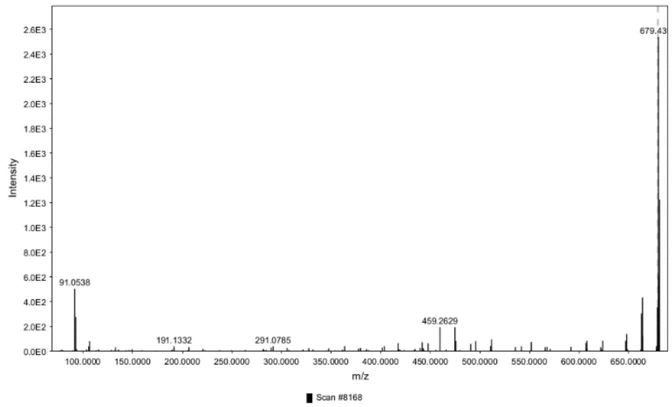


FigureS27

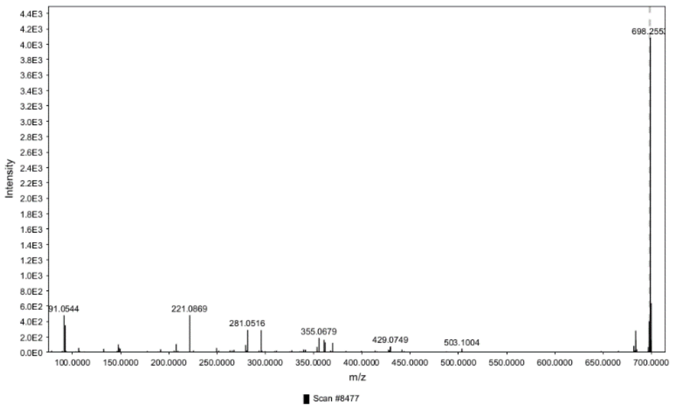


Figure S28

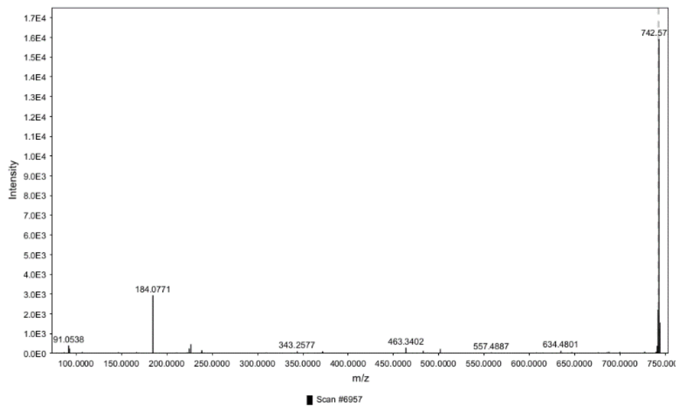


FigureS29

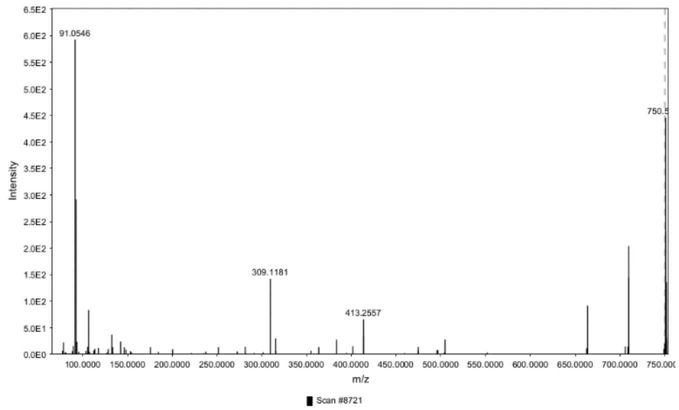


Figure S30

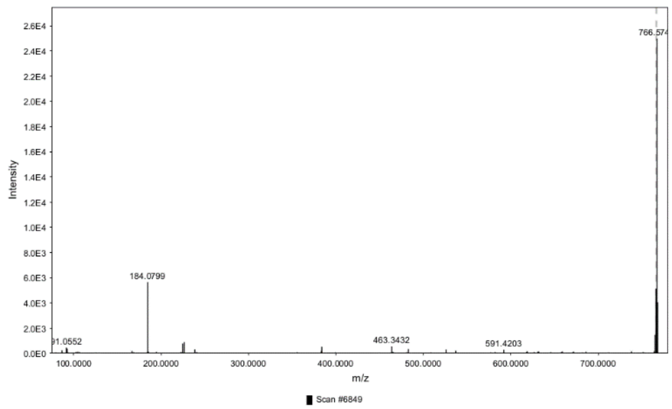


FigureS31

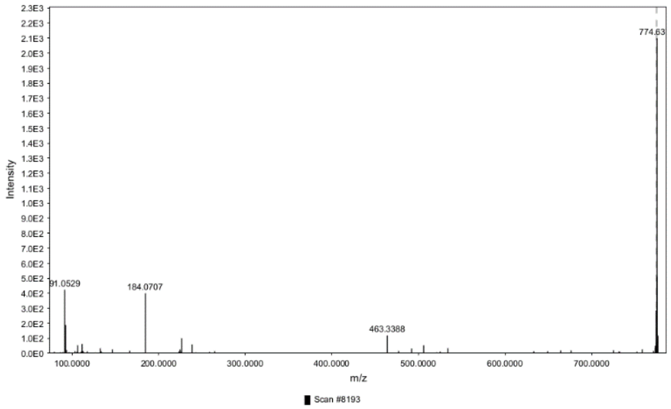


Figure S32

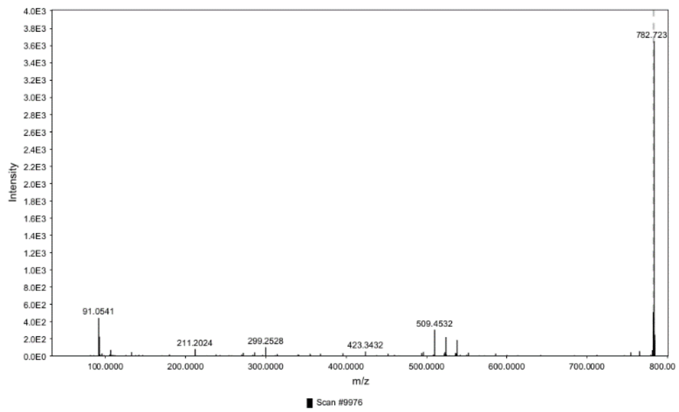


FigureS33

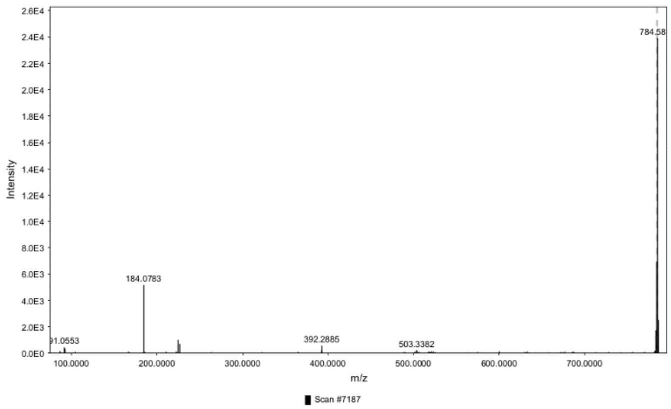


Figure S34

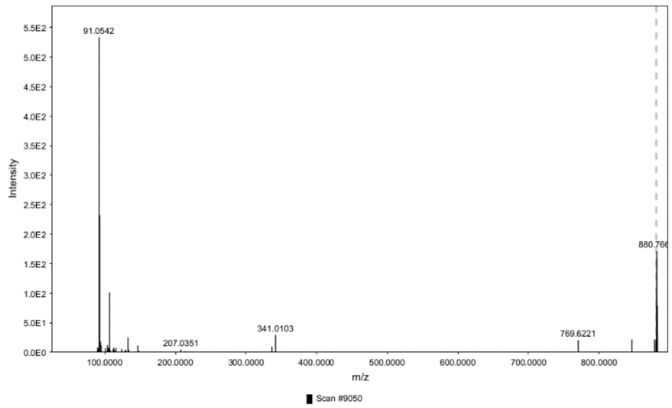


FigureS35

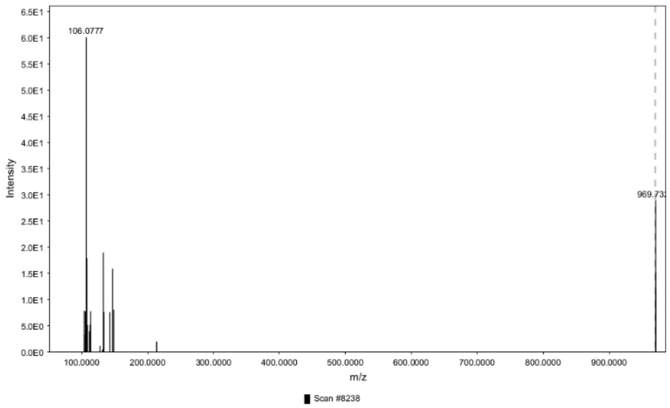


Figure S36

**Table S1: Zeno-MRM$^{HR}$-EAD parameters in HILIC(+) for selected features exhibiting a VIP above 1.5.**

| Retention time (min) | Precursor *m/z* | TOF MS/MS start *m/z* | TOF MS/MS stop *m/z* | Accumulation time (ms) | DP (V) | CE (V) | KE (eV) | Reaction time (ms) |
|---|---|---|---|---|---|---|---|---|
| 15.7 | 202.0690 | 30 | 207 | 110 | 60 | 15 | 10 | 20 |
| 10.3 | 204.0875 | 30 | 209 | 110 | 60 | 15 | 10 | 20 |
| 5.32 | 217.0680 | 30 | 222 | 110 | 60 | 15 | 10 | 20 |
| 10.3 | 244.0793 | 30 | 249 | 110 | 60 | 15 | 10 | 20 |
| 14.8 | 434.1649 | 44 | 439 | 110 | 60 | 15 | 10 | 20 |
| 14.0 | 442.2668 | 45 | 447 | 110 | 60 | 15 | 10 | 20 |
| 6.45 | 461.2764 | 47 | 466 | 110 | 60 | 15 | 10 | 20 |
| 14.8 | 549.2684 | 56 | 554 | 110 | 60 | 15 | 10 | 20 |

**Table S2: Zeno-MRM$^{HR}$-EAD parameters in RPLC(+) for selected features exhibiting a VIP above 1.5.**

| Retention time (min) | Precursor *m/z* | TOF MS/MS start *m/z* | TOF MS/MS stop *m/z* | Accumulation time (ms) | DP (V) | CE (V) | KE (eV) | Reaction time (ms) |
|---|---|---|---|---|---|---|---|---|
| 5.43 | 565.4033 | 57 | 570 | 50 | 60 | 15 | 15 | 20 |
| 4.98 | 284.2942 | 29 | 289 | 50 | 60 | 15 | 15 | 20 |
| 8.07 | 742.5756 | 75 | 748 | 50 | 60 | 15 | 15 | 20 |
| 5.41 | 605.3956 | 61 | 610 | 50 | 60 | 15 | 15 | 20 |
| 10.6 | 880.7608 | 89 | 886 | 50 | 60 | 15 | 15 | 20 |
| 9.65 | 550.5092 | 56 | 556 | 50 | 60 | 15 | 15 | 20 |
| 9.52 | 679.4302 | 68 | 684 | 50 | 60 | 15 | 15 | 20 |
| 9.66 | 548.5032 | 55 | 554 | 50 | 60 | 15 | 15 | 20 |
| 9.40 | 969.7362 | 98 | 975 | 50 | 60 | 15 | 15 | 20 |
| 9.91 | 698.2554 | 70 | 703 | 50 | 60 | 15 | 15 | 20 |
| 9.38 | 587.5485 | 59 | 593 | 50 | 60 | 15 | 15 | 20 |
| 8.24 | 784.5848 | 79 | 790 | 50 | 60 | 15 | 15 | 20 |
| 11.7 | 782.7221 | 79 | 788 | 50 | 60 | 15 | 15 | 20 |
| 8.01 | 766.5737 | 77 | 772 | 50 | 60 | 15 | 15 | 20 |
| 9.45 | 774.6373 | 78 | 780 | 50 | 60 | 15 | 15 | 20 |
| 10.2 | 750.4069 | 76 | 755 | 50 | 60 | 15 | 15 | 20 |

**Table S3: Filtering steps on the datasets.** For each step, a global percentage of removed features is calculated.

| **Chromatographic separation (polarity)** | **Number of input features** | **Blank filter** | **QC filter** | **Missing values filter** | **Univariate non-parametric test** | **Dead time filter** |
|---|---|---|---|---|---|---|
| HILIC(+) | 1964 | 1698 | 1460 | 1417 | 114 | 95 |
| HILIC(-) | 1488 | 1266 | 98 | 98 | 20 | 11 |
| RPLC(+) | 936 | 934 | 894 | 894 | 172 | 159 |
| RPLC(-) | 154 | 152 | 143 | 143 | 41 | 19 |
| **Step-by-step removed features percentage** | | 11% | 36% | < 2% | 86% | 18% |

**Table S4: Summary of the PLS-DA models performance parameters.**

| **Dataset** | **$R^2X$** | **$R^2Y$** | **$R^2Y$ p-value** | **$Q^2$** | **$R^2Y$ p-value** | **RMSEE*** |
|---|---|---|---|---|---|---|
| HILIC(+) | 0.685 | 0.938 | < 0.05 | 0.898 | < 0.05 | 0.133 |
| HILIC(-) | 0.676 | 0.902 | < 0.05 | 0.745 | < 0.05 | 0.171 |
| RPLC(+) | 0.712 | 0.709 | < 0.05 | 0.417 | < 0.05 | 0.288 |
| RPLC(-) | 0.620 | 0.773 | < 0.05 | 0.493 | < 0.05 | 0.255 |

*, RMSEE stands for Root Mean Square Error of Estimation
